# Establishment of a prognosis prediction model based on pyroptosis-related signatures associated with the immune microenvironment and molecular heterogeneity in clear cell renal carcinoma

**DOI:** 10.1101/2021.08.05.455284

**Authors:** Aimin Jiang, Jialin Meng, Yewei Bao, Anbang Wang, Wenliang Gong, Xinxin Gan, Jie Wang, Yi Bao, Zhenjie Wu, Juan Lu, Bing Liu, Linhui Wang

## Abstract

**Background:** Pytoptosis is essential for tumorigenesis and progression of clear cell renal cell carcinoma (ccRCC). However, the heterogeneity of pyroposis and its relationship with the tumor microenvironment (TME) remain unclear. The aim of the present study was to identify proptosis-related subtypes and construct a prognosis prediction model based on pyroptosis signatures.

**Methods:** First, heterogenous pyroptosis subgroups were explored based on 33 pyroptosis-related genes and ccRCC samples from TCGA, and the model establsihed by LASSO regression was verified by ICGC database. Then, the clinical significance, functional status, immune infiltration, cell-cell communication, genomic alteration and drug sensitivity of different subgroups were further analyzed. Finally, the LASSO-Cox algorithm was applied to narrow down the candidate genes to develop a robust and concise prognostic model.

**Results:** Two heterogenous pyroptosis subgroups were identified: pyroptosis-low immunity-low C1 subtype, and pyroptosis-high immunity-high C2 subtype. Compared with C1, C2 was associated with a higher clinical stage or grade and a worse prognosis. More immune cell infiltration was observed in C2 than that in C1, while the response rate in C2 subgroup was lower than that in C1 subgroup. Pyroptosis related genes were mainly expressed in myeloid cells, and T cells and epithelial cells might influence other cell clusters via Pyroptosis related pathway. In addition, C1 was characterized by MTOR and ATM mutation, while C2 was characterized by more significant alterations in SPEN and ROS1 mutation. Finally, we constructed and validated a robust and promising signature based on the pyroptosis-related risk score for assessing the prognosis in ccRCC.

**Conclusion:** We identified two heterogeneous pyroptosis subtypes and 5 reliable risk signatures to establish a prognosis prediction model. Our findings may help better understand the role of pyroptosis in ccRCC progression and provide a new perspective in the management of ccRCC patients.

## Introduction

Renal cell carcinoma (RCC) is a common urologic malignancy with an incidence only secondary to prostate and bladder cancer^1^. According to the characteristics of molecular biology and histopathology, RCC can be categorized into two main types: clear cell renal cell carcinoma (ccRCC), and non-clear cell renal carcinoma (nccRCC) including papillary renal cell carcinoma (pRCC), chromophobe cell renal cell carcinoma (cRCC) and collecting duct renal cell carcinoma (cdRCC)^2^. Among them, ccRCC accounts for approximately 75-80%. As only 6-10% patients developed typical symptoms like backache, an abdominal mass or hematuria, it is difficult to diagnose RCC in the early stage^3^. As ccRCC is insensitive to conventional chemotherapy and radiotherapy, nephrectomy, target therapy and immunotherapy are the mainstay of treatment for ccRCC^4^. But as ccRCC is an extremely heterogeneous disease, even patients with similar clinical characteristics wo received similar treatments may have distinctive outcomes^5^. Hence, it is urgent to explore the innate mechanism of ccRCC for the sake of developing novel therapeutic strategies for improving the overall clinical outcome of this disease.

Cell death is not only a physiological regulator of cell proliferation, stress response and homeostasis but a tumor inhibitive mechanism^6^. There are several known cell death types, including necrosis, apoptosis, necroptosis, autophagy, anoikis and pyroptosis. Apoptosis has been extensively and thoroughly investigated as a significant mechanism of anti-cancer defense, but the relationship between pyroptosis and cancer remains unclear. Pyroptosis is an inflammatory form of cell death triggered by certain inflammasomes, leading to the cleavage of gasdermin D (GSDMD) and activation of inactive cytokines interleukin-18 (IL-18) and IL-1β. Recently, extensive studies have focused on elucidating the molecular mechanism underlying pyroptosis as well as the mechanism of inducing pyroptosis in tumor cells^7^. Tan et al reported that BRD4 inhibition prevented RCC cell proliferation and epithelia-mesenchymal transition (EMT) progression, and exerted an antitumor effect in RCC by activating the NF-κB-NLRP3-caspase-1 pyroptosis signaling pathway^8^. All these findings suggest that pyroptosis play an essential role in the progression and therapy in ccRCC, and comprehensive analysis of pyroptosis may shed new light on the development of strategies for the treatment of ccRCC. However, the accurate mechanism of pryoptosis in ccRCC has been less studied. Herein, we aimed to perform a systematic research to compare the expression level in ccRCC and normal renal tissue, decipher the role of pyroptosis in the ccRCC microenvironment and construct a pyroptosis related risk model for ccRCC.

## Materials and Methods

### Public dataset collection

ccRCC data were enrolled from The Cancer Genome Atlas (TCGA) cohorts(n=607) and International Cancer Genome Consortium(ICGC) cohorts(n=91). For datasets in the TCGA and ICGA databases, institutional review board approval and informed consent were not required. Level-3 transcriptome and clinical information were download form TCGA and ICGC. Patients were excluded if they 1) did not have prognostic information, and 2) died within 30 days. The overall workflow of this study is displayed in **Figure S1**.

### Identification of differentially expressed genes (DEGs) related to pyroptosis

Altogether 33 pyroptosis-related genes were retrieved from prior articles and reviews (**Supplement Table S1**). Correlations between these pyroptosis-related genes were assessed by Spearman’s rank correlation using R ‘corrplot’ package. The cluster analysis of pyroptosis-related genes was performed by hclust and kmeans algorithms. Then, 531 ccRCC patients were categorized into different subgroups using PCA, and finally the subtype number k = 2 was selected in that it turned out to be the best classifier number. R package ‘DEseq2’ was employed to identify DEGs between different groups, with the threshold set as *p*-adjusted value < 0.01 and abstract log-foldchange = 2. To explore the potential molecular mechanisms underlying the subgroups, R package ‘clusterProfiler’ was used to perform gene ontology (GO), Kyoto Encyclopedia of Genes and Genomes (KEGG) pathway and Gens set enrichment analysis (GSEA). Gene set permutations were performed 1000 times for each analysis. Gene sets with FDR < 0.01 were considered significantly enriched.

### Classification of the pyroptosis status

Based on 33 pyroptosis gene expression matrix data sets of 516 ccRCC patients, two different pyroptosis status groups (C1 and C2) were identified by using R ‘ConseensusClusterPlus’ package. Principal component analysis (PCA) was analyzed and visualized by R ‘ggord’ and ‘ggplot’ packages.

### Analysis of the DEGs

DEseq2 package was used to identify DEGs between C1 and C2, with the adjusted P value<0.05 and abstract logFC>1.2 considered a significant difference. Then Gene Ontology (GO) enrichment analysis including biological process (BP), cellular components (CC) and molecular function(MF). The Cytoscape plugin iREgulon was employed to ananlyze transcription of the down- and up-regulated genes, and the iRegulon plugin could identify regulons using motifs and track discovery in an existing network or in a set of coregulated genes. KEGG and GSEA analyses were performed by using R ‘clusterprofiler’ packages, and the difference in signaling pathways between C1 and C2 subgroups were presented by adoption of the gene set from MSigDB database^9^. The potential transcription factors for DEGs were analyzed by using module ‘iRegulon’ form Cytoscape software.

### Differences in tumor microenvironment (TME) and immunotherapy response

To quantify the proportion of immune cells between the subtypes, several immune-related algorithms including TIMER, CIBERSORT, QUANTISEQ, MCPCOUNTER, XCELL and EPIC were employed to calculate the cellular components or immune cell enrichment scores in ccRCC tissues, and differences between C1 and C2 subgroups were compared. Single sample gene set enrichment analysis (ssGSEA) was employed to quantify the relative abundance of 28 immune cells in ccRCC TME^10–13^. Differences in immune cell infiltration in TME were visualized by Heatmap and boxplot. R ESIMATE package was used to identify the stromal component and immune component between the two subgroups. Tumor Immune Dysfunction and Exclusion (TIDE) algorithms were applied to predict the immunotherapy response of each ccRCC patient.

### Cell-cell interaction analysis

PRJNA705464, which is a large database containing cells totally, was applied to investigate the role of pyroptosis-related genes in cell-cell interaction in ccRCC TME. Only untreated tumor samples from PRJNA705464 were selected for further analysis. R package “Seurat” was applied for dimension reduction and clustering analysis, and R package “SingleR” was introduced for cell type identification^14^. R package “CellChat” and software “cellphonedb” were applied for cell-cell interaction analysis, and cell–cell interactions based on the expression of known L–R pairs in different cell types were calculated^15, 16^. In brief, gene expression data of cells and assigned cell types were used as input for CellChat. Firstly, overexpressed ligands or receptors in one cell group were identified, and then gene expression data were projected onto the protein-protein interaction network. The overexpressed L–R interactions were identified if either the ligand or receptor was overexpressed. Next, CellChat was used to infer the biologically significant cell–cell communication by assigning each interaction with a probability value and performing a permutation test. Finally, communication networks were visualized using circle plot and signaling pathways visualized using bubble plot.

### Multi-omics data analysis

Mutation and copy number variations were downloaded from TCGA database. WES data were used to compare differences in somatic mutation between C1 and C2 using the R ‘maftools’ package^17^. Fisher’s exact test was introduced to identify the different mutation genes with a *P* value <0.05. The co-occurrence and mutually exclusive mutation were identified using the CoMEt algorithms. For copy number variation data, th GISTIC 2 software in GenePatterns was applied to identify significantly deleted or amplified broad and focal segments^18^.

### Construction and validation of the pyroptosis-related risk scores in public data sets

To assess the prognostic value of the pyroptosis-related genes, a single analysis was first performed, and 17 survival-related genes were selected for further analysis. Then, lasso-cox algorithm was applied to narrow down the candidate genes to develop a robust and concise prognostic model. Ultimately, a risk model based on five genes was constructed and the penalty parameter was decided by 1se using R ‘glmnet’ package. After standardization and normalization of the TCGA ccRCC expression data, the risk score of each patient was calculated using the following equation: Risk score = *0.0271113*AIM2+0.04147645*GSDMB+0.01748664*IL6+0.01968024*PYCARD-0.08678271*TIRA P* 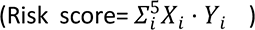. Then, each patient from TCGA and ICGC database was assigned to either a high-risk group or a low-risk group based on the median value of the risk score. Kaplan-Meier survival curves were depicted to predict the clinical outcomes in the two groups using R ‘survival’ package. Differences in survival between the two groups were evaluated by log-rank test. The ROC curves were depicted and the area under the curves (AUC) for 0.5-,1-, 2-, 3- and 5-year overall survival (OS) and progression-free interval (PFI) were calculated using R ‘timeROC’ package.

### Assessment of clinical significance of the pyroptosis subtypes

Clinical characteristics including age, gender, grade, AJCC stage, TNM, OS and PFI were compared between C1 and C2 subtypes by R ‘compare’ package. Sensitivity to several chemotherapy drugs were compared by R ‘pRRohetic’ package^19^. IC50(half maximal inhibitory concentration) values of C1 and C2 subgroups were estimated by ridge regression. The sensitivity of the two subgroups to immune check point inhibitor therapy was predicted by TIDE (http://tide.dfci.harvard.edu) algorithm. The Genomics of Drug Sensitivity in Cancer (GDSC) database (https://www.cancerrxgene.org) was applied to screen the potential drug for the high-risk subgroup by R ‘pRRphetic’ package^20^.

### Validation of risk model-related gene expression in the SMMU cohort

According to the expression of prognostic genes in the gene signature in TCGA and ICGC database, we selected five hub genes (PYCARD, AIM2, IL6, GSDMB and TIRAP) that were differentially expressed between the cancer and normal tissues for validation using quantitative real-time PCR (RT-qPCR). Informed consent about the tissue sample analysis was obtained from each patient before initiation of the study, and the study protocol was approved by the Institutional Review Board of the Second Military Medical University (SMMU) Cancer Center. A total of 40 pairs of normal and cancer tissues were stored at −80 ℃ before use. Total RNA was extracted from the tissue samples using TRIzol reagent (Thermo Fisher Scientific, Waltham, MA, USA). The concentration of the isolated RNA was measured with the NanoDrop2000 Spectrophotometer (Thermo Scientific, Wilmington, DE, USA). Total RNA (2 μg) was reverse-transcribed and RT-qPCR was carried out on the triplicate samples in an SYBR Green reaction mix (Takara Biotechnology, Shiga, Japan) with an ABI Quant Studio5 Real-Time PCR System (Applied Biosystems, Carlsbad, CA, USA). The primer sequences used are listed in **Table S1** (see Supplementary Data). Glyceraldehyde 3-phosphate dehydrogenase (GAPDH) was employed for normalization. Reactions without an RNA template or reverse transcriptase were used as negative controls. The expression of individual RNA molecules was determined by the -ΔCT approach (ΔCT=CT _RNA_ - CT _GAPDH_RNA_). All procedures for RT-qPCR were performed according to the manufacturer’s protocol.

### Statistical analysis

Differences in the expression of the pyroptosis-related genes in the public data sets were compared by One-way ANOVA, and differences in clinical information and immune check point inhibitor response between the two different subgroups were compared by Chi-squared test. Differences in OS and PFI between the two subgroups were compared by Kaplan-Meier method and log-rank rest. The hazard ratios (HRs) were calculated by univariate Cox regression and multiple Cox regression analysis. The receiver operating characteristic (ROC) curves were plotted by ‘timeROC’ R package. The performance of the risk score in predicting OS and PFI was evaluated by area under the ROC curve (AUC) and Harrell’s concordance index (C-index). All P-values were two-sided, with P < 0.05 as statistically significant. Adjusted P-value was obtained by Benjamini-Hochberg (BH) multiple test correction. All data processing, statistical analysis and plotting were conducted using R 4.0.4 software.

## Results

### Landscape of pyroptosis genes in ccRCC

Firstly, the expression levels of 33 pyroptosis genes were compared between the ccRCC and normal renal tissues in TCGA dataset. The result showed that the expression level of NLRP1, NOD1, PLCG1, PLCG1, GSDMB, NLRP6, GSDMC, NLRP7, IL1B, GSDMA, CASP3, NLRRC4MNLRP3, CASP8, CASP1, CASP4, CASP5, AIM2, NOD2, GPX4, GSDMD, PYCARD and IL18 in the ccRCC tissues was higher than that in the normal renal tissues, while the expression level of genes containing CASP9 and NLRP2 in the normal renal tissues was higher than that in the ccRCC tissues (Figure 1A). To further explore the interaction and correlation of the pyroptosis genes, we constructed a comprehensive network and divided the genes into 4 clusters. It was found that three pyroptosis genes were risk prognosis factors (Figure 1B).

**Figure 1.**
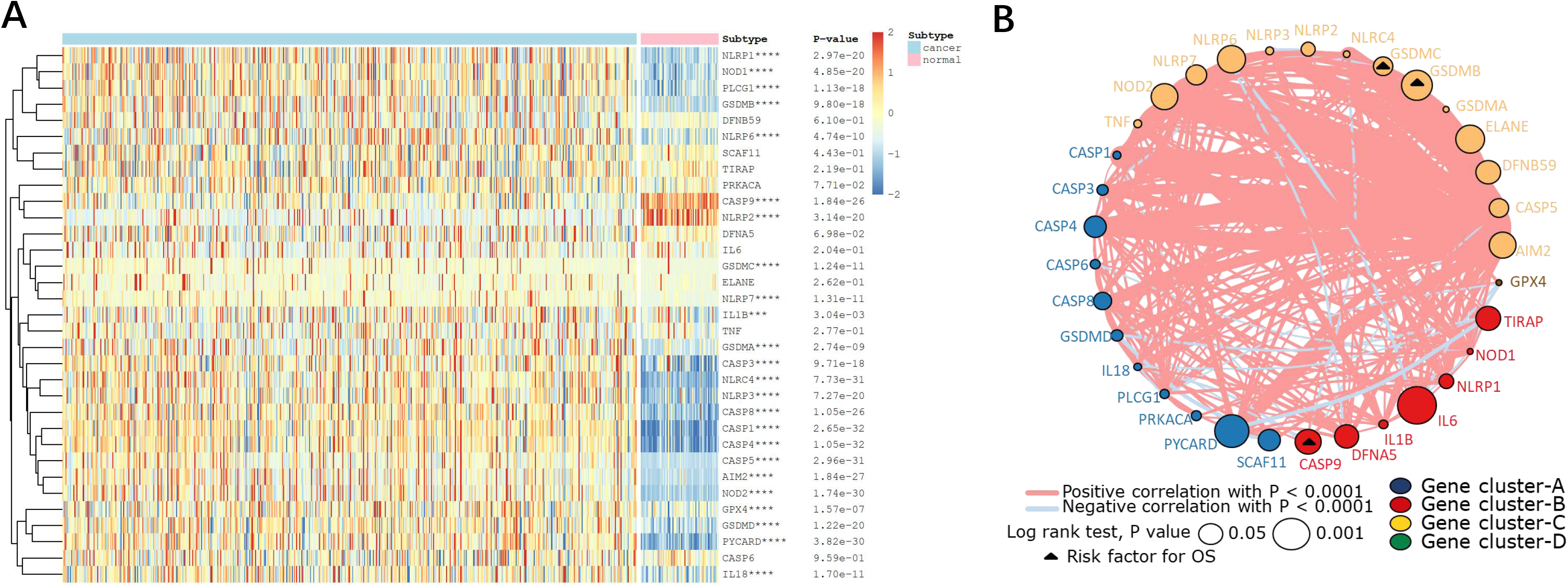
The landscape of pyroptosis-related genes in ccRCC. (A) The expression levels of 33 pyroptosis-related genes in ccRCC. The darker color indicates a higher expression, where the red color indicates up-regulation and the blue color indicates down-regulation. The upper tree diagram represents clustering results for different samples from different experimental groups, and the left tree shows the results of cluster analysis for different genes from different samples. (B) Interaction of pyroptosis-related genes. Gene cluster A, blue; gene cluster B, red; gene cluster C, yellow; gene cluster D, green. The circle size represents the effect of each gene on prognosis, and the range of values was calculated by log-rank test as p<0.05 and p<0.001. The triangle dots in the circle represent the risk factors of prognosis. The lines linking regulators show their interactions, and thickness shows the correlation strength between the regulators. A negative correlation is marked with blue and a positive correlation with red.

### Two clusters of ccRCC are identified by consensus clustering of pyroptosis genes

After removing the normal renal tissues, we used unsuperbised clustering methods to classify the tumor samples into different molecular subgroups based on pyroptosis-related genes. The optimal cluster number was identified by R ‘ConsensusClusterPlus’ package, and the clustering stability was evaluated by the proportion of the PAC algorithm. Finally, two distinct clusters, termed as C1 and C2, were identified (Figure 2A-C). To better understand the clustering result, clinical outcomes and clinicopathological features, differences in survival in terms of OS and PFI were compared between the two clusters by log-rank test and Kaplan-Meier curve (Figure 2D-E). In addition, we found that most pyroptosis-related genes were highly expressed in C2 as compared with C1 (Figure 2F). Compared with C1, C2 was significantly correlated with a higher grade, AJCC score and TNM status (Table 1).

**Figure 2.**
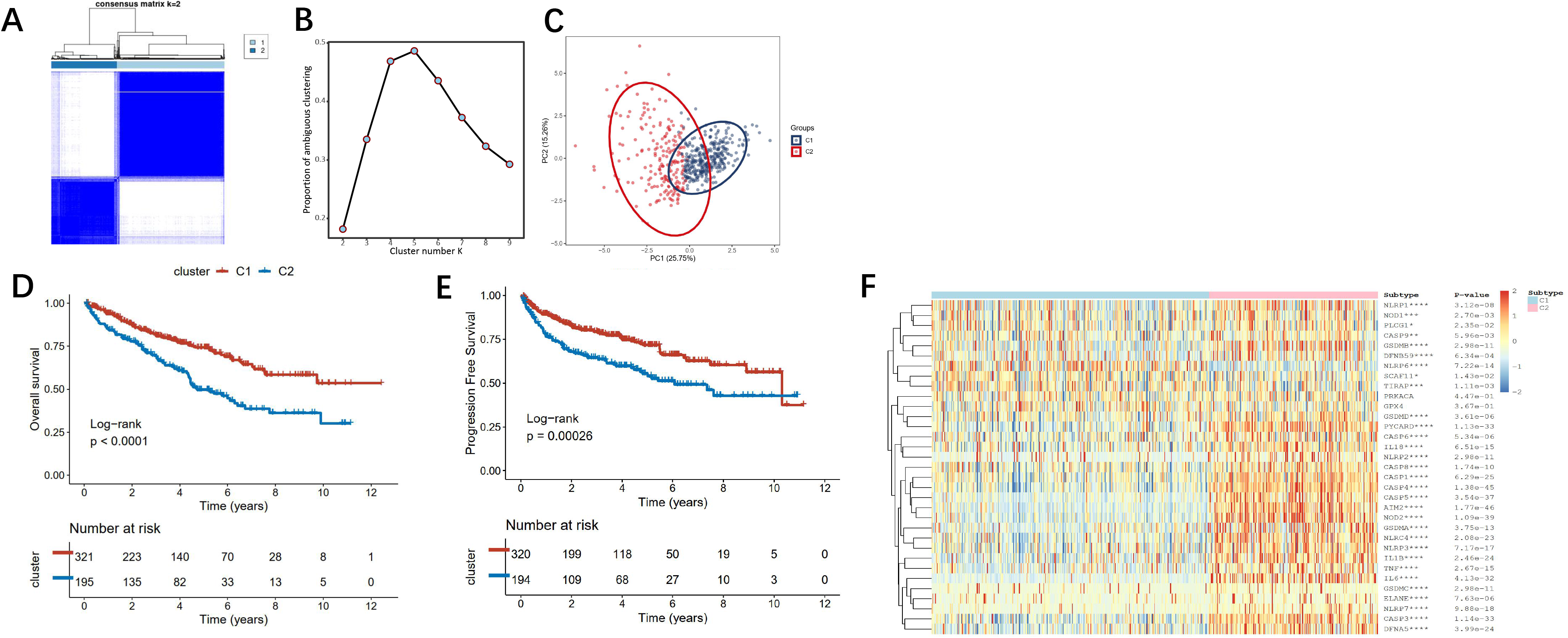
Identification of pyroptosis clusters of ccRCC. (A) The consensus score matrix of all samples when k = 2. A higher consensus score between two samples indicates that they are more likely to be grouped into the same cluster in different iterations. (B) The proportion of ambiguous clustering (PAC) score, where a low value of PAC implies a flat middle segment, allowing conjecture of the optimal k (k = 2) by the lowest PAC. (C) Two-dimensional principal component plot by the expression of 33 pyroptosis-related genes in the two subtypes. The blue dots represent C1, and red dots represent C2. (D-E) Kaplan-Meier analysis for overall survival (left) and progression-free interval of the two subtypes in the TCGA cohort; (E) The expression heatmap of the 33 pyroptosis-related genes in the two subtypes.

**Table 1.**
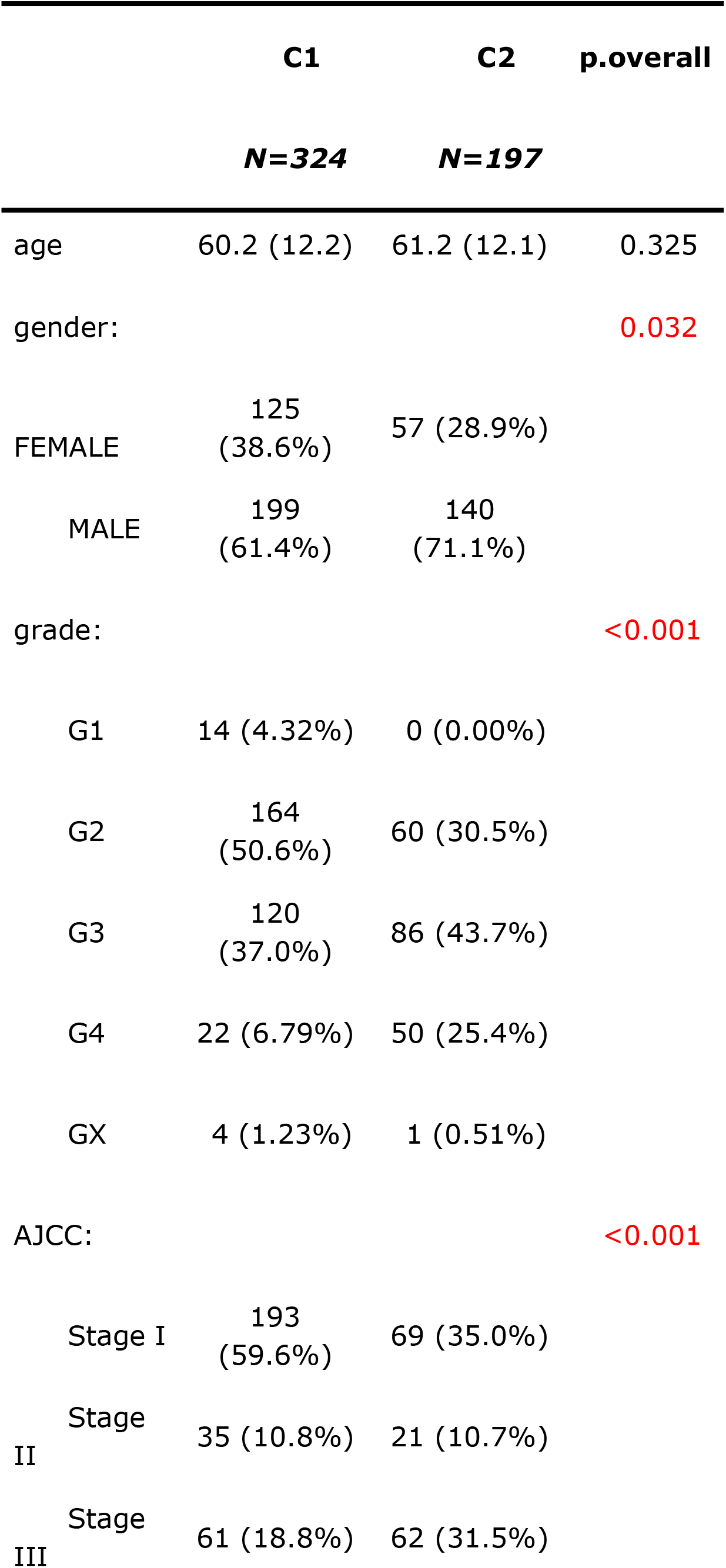
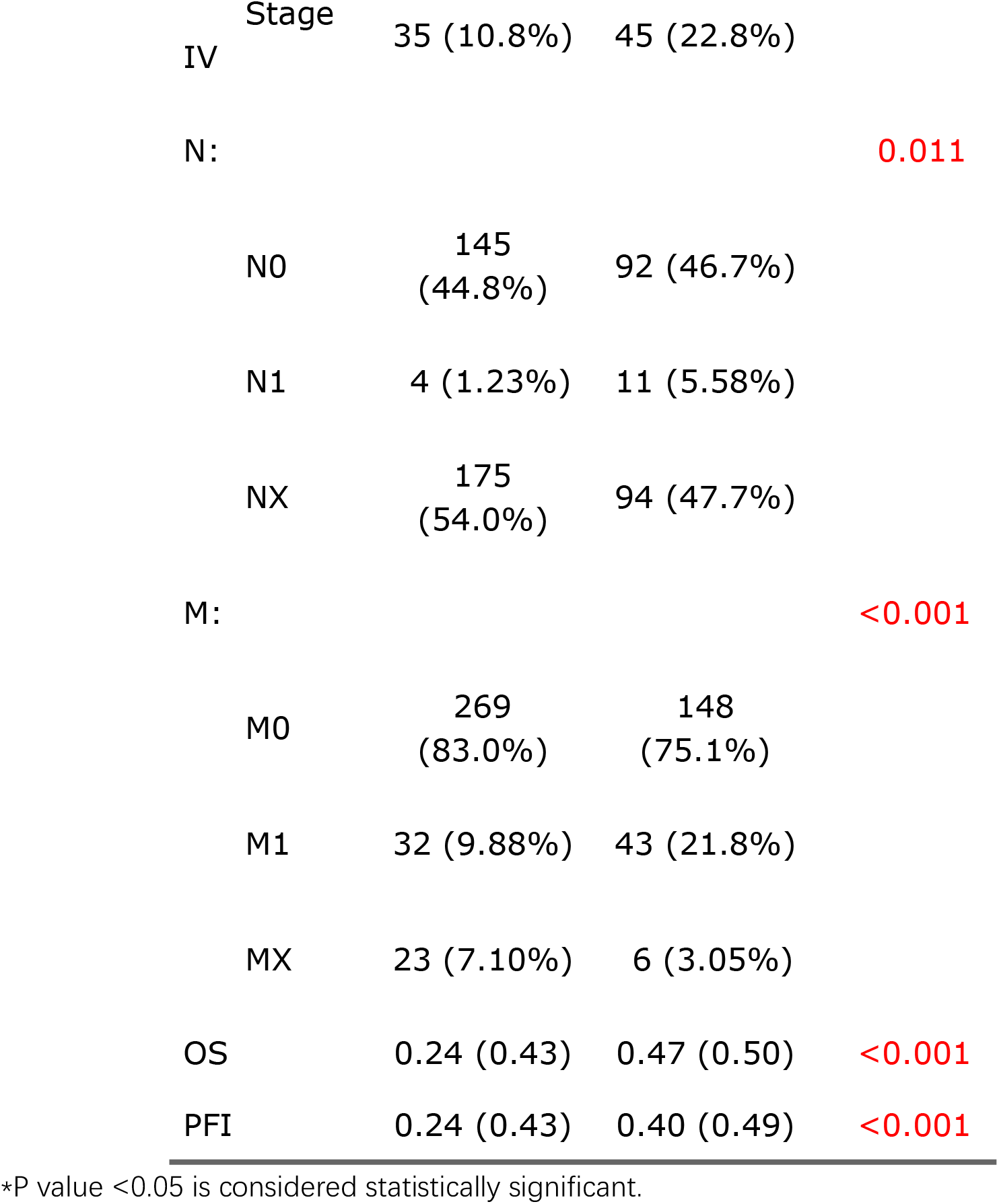
Difference of clinical characteristics between C1 and C2 subgroups

### Identification of DEGs and functional analysis

The gene expression profiles of ccRCC were analyzed to identify pyroptosis-related DEGs, including the up-regulated and down-regulated ones in C2 relative to C1 (Figure 3A). Then, DEGs were used to perform GO enrichment, KEGG pathway, GSEA and GSVA analyses. The GO results demonstrated that the DEGs were enriched in humoral immune response, receptor ligand activity and collagen-containing extracellular matrix (Figure 3B**, S2A**). Transcription factor analysis of the downregulated and unregulated genes were conducted using iRegulon, a Cytoscape plugin, and a normalized enrichment score (NES) >10 was considered to be significant. The transcriptional regulation network of these down- and up-regulated genes were shown in Figure S. GSEA analysis showed that the adaptive immune system, cytokine signaling in the immune system and hemostasis were upregulated, while eukaryotic translation termination, peptide chain elongation and regulation of apoptosis were downregulated in C2 *vs.* C1(Figure 3C-D). The TF was .GSVA analysis indicated that fatty acid metabolism, adipogenesis and PI3K-Akt-mtor pathway were upregulated in C2, while inflammatory response, apoptosis and IL6-JAK-STAT3 pathway were upregulated in C1(Figure 3E). The KEGG results demonstrated that cytokine receptor interaction and primary immunodeficiency were upregulated, while collecting duct acid secretion was down-regulated in C2 relative to C1(Figure 3F).

**Figure 3.**
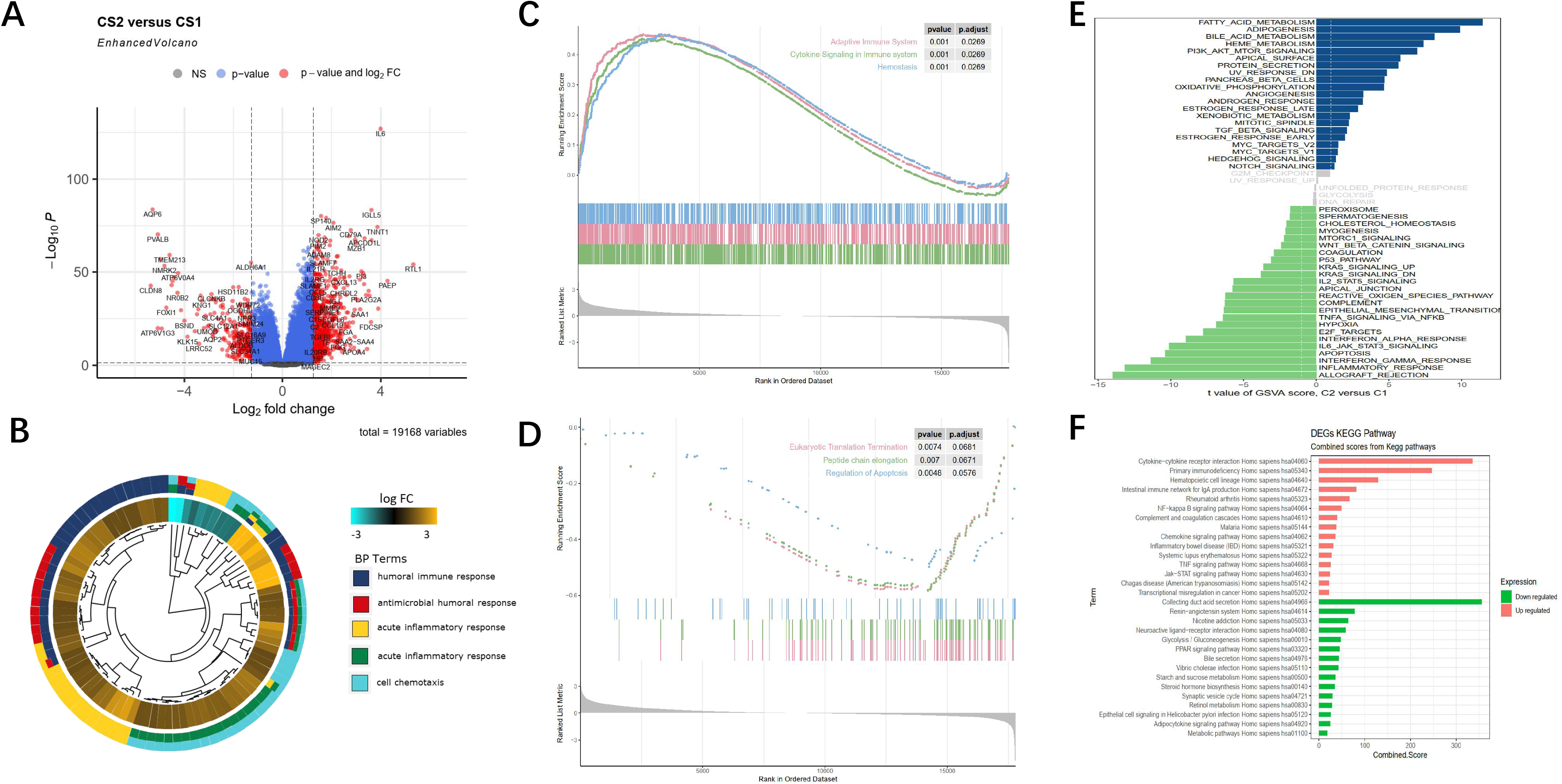
Functional enrichment analysis of DEGs between C1 and C2 subtypes. (A) Volcano map of differentially expressed genes. (B) GO enrichment analysis. (C-D) GSEA analysis shows the hallmarks between the subgroups. (E) Gene set enrichment analysis (GSVA) shows the significant enrichment differences between the subgroups. (F) KEGG pathway analysis.

### Comparison of the immune landscape between the subgroups

The heatmap of immune response based on different immune-related algorithms (including TIMER, CIBERSORT, CIBERSORT-ABS, QUANTISEQ, XCELL and EPIC) is depicted in Figure 4A. Then, single-sample gene set enrichment analysis (ssGSEA) was introduced to compare the immune cell enrichment scores between C1 and C2(Figure 4B). It was found that most immune cells, including activated B, CD4 T, CD8 T and dendritic cells, CD56 bright natural killer cell, central memory CD4 T cell, central memory CD8 T cell, effector memory CD4 T cell, effector memory CD8 T cell, eosinophil, Gamma delta T cell, immature dendritic cell, macrophage, mast cell, MDSC, memory B cell, monocyte, natural killer cell, natural killer T cell, regulator T cell, T follicular helper cell, type 1 T helper cell, type 17 T helper cell and type 2 T helper cell were all highly infiltrated in C2 subgroup. Only neutrophil was highly infiltrated in C1 subgroup. Then, the expression level of 9 immune check inhibitor genes was compared between C1 and C2 subgroups. It was found that most of these genes (including CD274, CD276, CTLA4, CXCR4, IL6, LAG3, PDCD1 and TGFB1) were upregulated in C2 subgroup (Figure 4C). Meanwhile, R ‘estimate’ package was utilized to investigate the immune-related scores between C1 and C2, and all the immune-related scores (including stromal score, immune score and estimate score) were significantly higher in C2 subgroup (Figure 4D). R ‘GSVA’ package was used to compare the immune-related signal enrichment scores between C1 and C2, and the result was consistent with the result forehead, indicating that all immune signals were more highly enriched in C2 (Figure 4E). Finally, by using TIDE algorithm, we compared the sensitivity of the immune check point inhibitors between the two subgroups, and found that the response rate in C1 subgroup was higher than that in C2 subgroup (40.8% *vs.* 24.6%) (Figure 4F).

**Figure 4.**
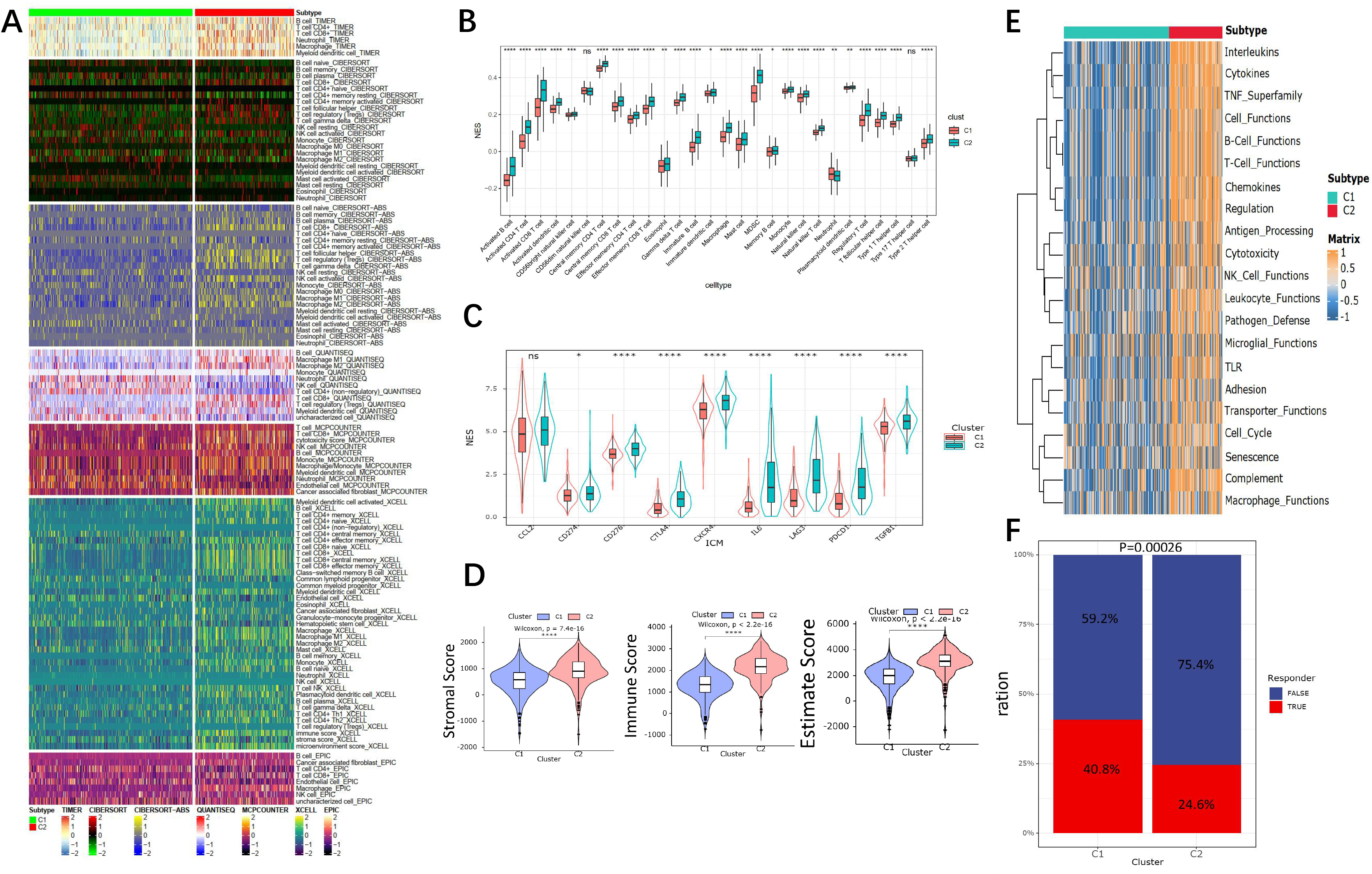
Immune landscapes between the pyroptosis subgroups. (A) Heatmap of tumor related infiltrating immune cells based on TIMER, CIBERSORT, CIBERSORT-ABS, QUANTISEQ, MCPcounter, XCELL and EPIC algorithms between the subgroups. (B) Different normalized enrichment scores of immune cells between the subgroups. (C) Different expressions of the immune check point inhibitor between the subgroups. (D) Differences in ESTIMATE score between the subgroups. (E) Heatmap of different immune related pathway enrichment scores between the subgroups. (F) Differences in response to the immune check point inhibitor treatment based on TIDE algorithm.

### Crosstalk between cancer and immune cells based on pyroptosis

To identify the role of pyroptosis-related genes in the TME of ccRCC, we collected the single cell sequence datasets from ccRCC patients who had never received any drug therapy, totally containing 29799 cells. We next used nonlinear dimensionality reduction (t-distributed stochastic neighbor embedding, t-SNE) and graph-based Louvain clustering algorithm to investigate cell distribution and heterogeneity of ccRCC, which included 1252 endothelial cells, 5552 epithelial cell,1329 mast cells, 7151 myeloid cells, 81 naive B cells, 152 plasma cells, 192 smother muscle cells, 11909 T cells, and 2142 unknow cells (Figure 5A). The expression level of pyroptosis-related genes in myeloid cells was significantly higher than that in other cells (P < 0.01) (Figure 5B). To investigate the impact of pyroptosis-related genes on cell-cell communication, we used ‘CellChat’ and “cellphonedb” to analyze the crosstalk between cancer and immune cells, and determine the complex cell-cell interaction network between cancer and immune cells (Figure 5C-D). All the cell-cell communications among cells were explored via CellChat and cellphonedb,respectively(**Figure S3**). Next, we explored the pathways involved in proptosis and found that TNF pathway (including TNFRSF1B-GRN and TNFRSF1A-GRN) could trigger pyroptosis in myeloid cells (Figure 5E-G). Our correlation analysis further verified the above results (Figure 5H). In summary, our results revealed that pyroptosis-related genes had the potential to shape the unique TME of ccRCC.

**Figure 5.**
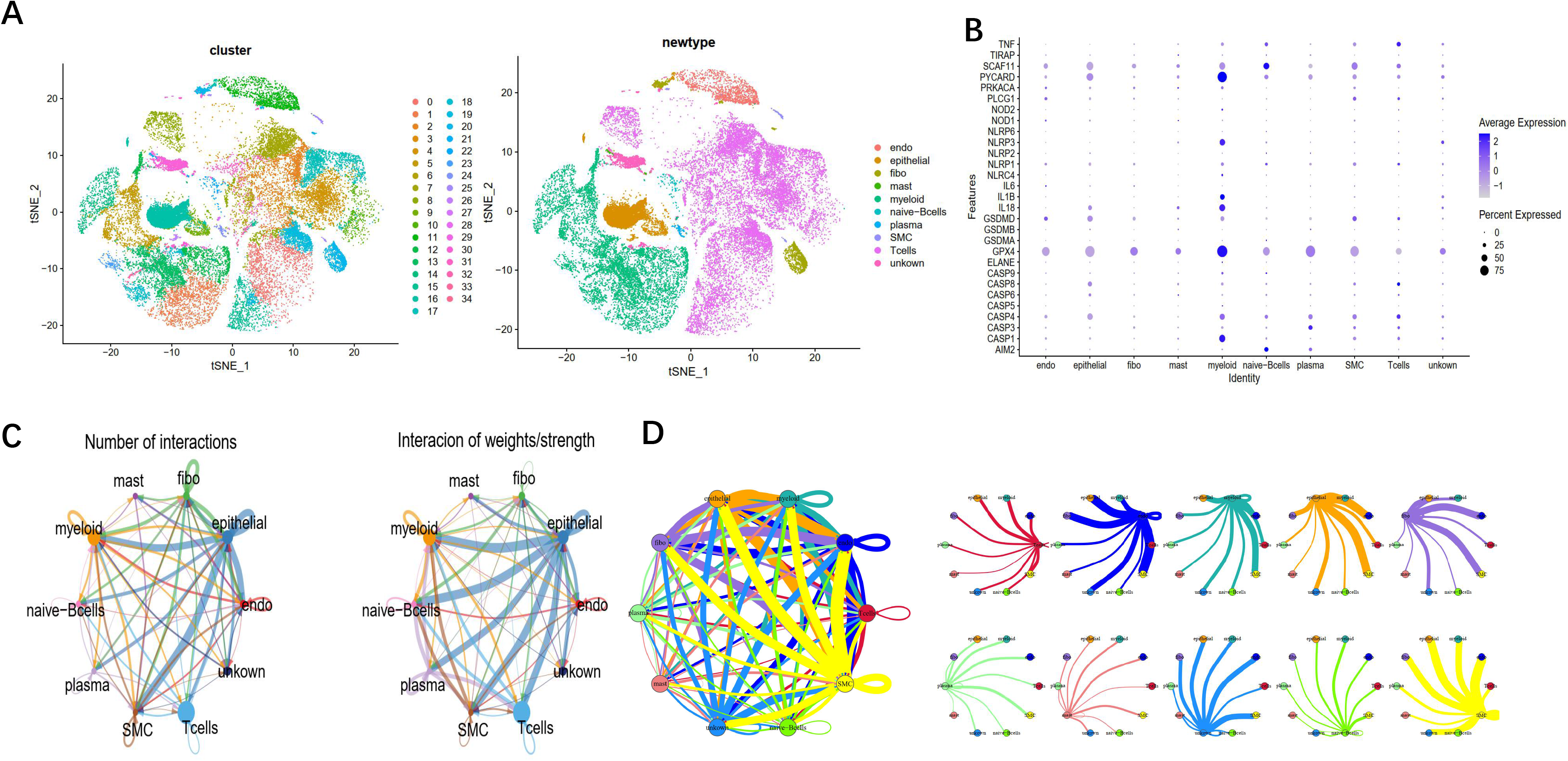
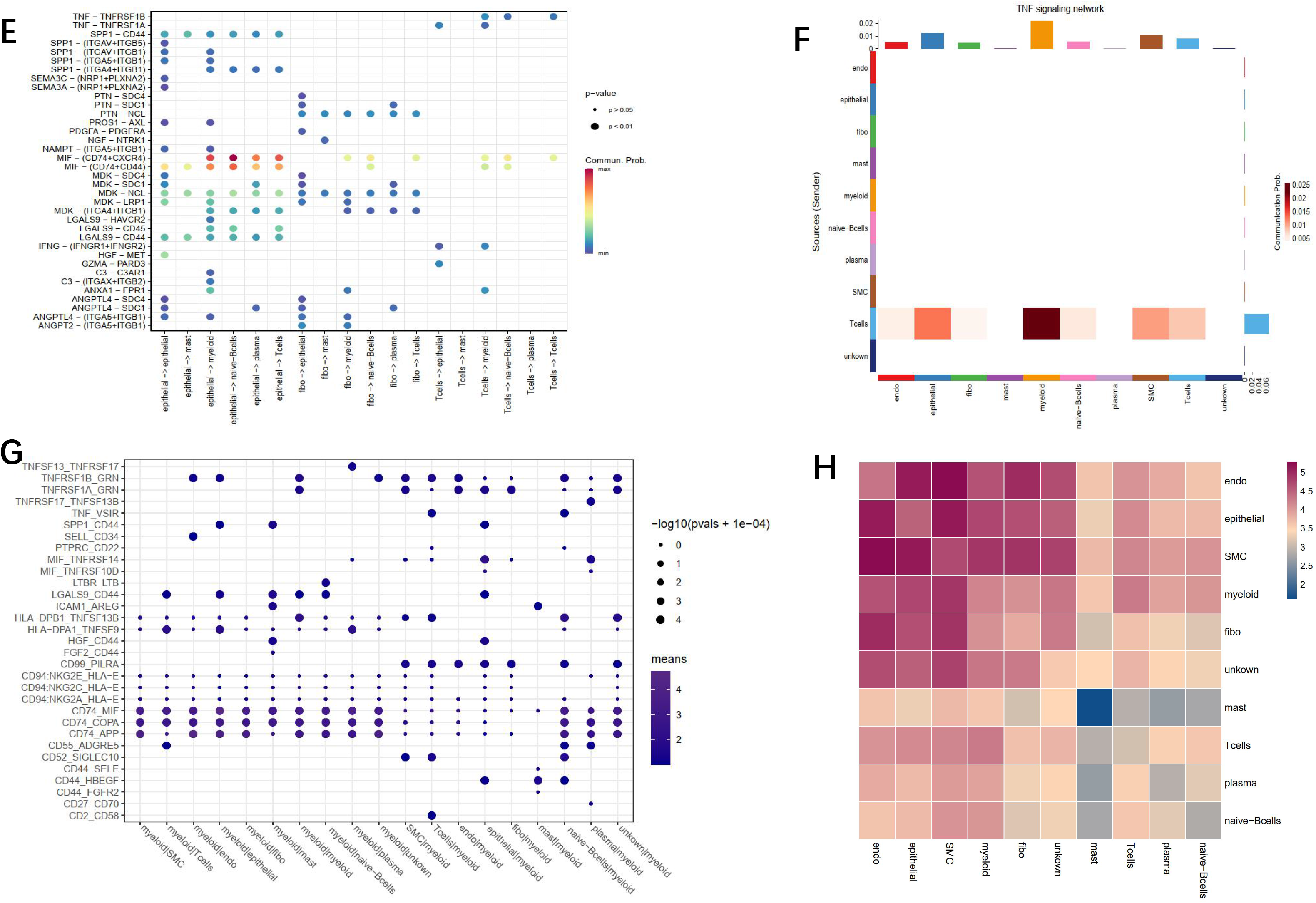
Crosstalk between cancer and immune cells. (A) The t-SNE plot shows that 29799 cells were divided into 35 clusters, among which T cell accounts for the largest cluster. (B) Pyroptosis-related genes expressed in the 10 cell clusters. (C) Cell-cell interaction in cell clusters as analyzed by “CellChat”. (D) Cell-cell interaction in cell clusters as analyzed by “cellphonedb”. (E) Connection probability of main signaling pathways in cell clusters as analyzed by “CellChat”. (E) TNF signaling pathways between T cells and other cells. (G) Connection probability of main signaling pathways in cell clusters as analyzed by “cellphonedb”. (H) Correlation of the communication ratio between cell clusters.

### Comprehensive and integrated genomic characterization of the two subgroups

After detecting the transcriptional alterations in the above section, we further investigated the disparity in the genomic layer in the wo subgroups. It was fund that the mutation rate was similar between the two subgroups (185/214, 86.45% in C1 *vs.* 82/106, 77.36%). The top 20 most frequently mutated genes in the corresponding cohort are depicted in Figure 6A-B, showing significant differences between the two subgroups. We found that most of mutation genes were identified as protective factors in C2 subgroup(Figure 6C). Also, the tumor mutation burden rate in C2 was higher than that in C1 (Figure 6D). Next, the co-occurrence and exclusive mutations of the top 20 most frequently mutated genes were investigated by using CoMEt algorithm. Compared with the pervasive co-occurrence landscape, there were unique cases in the two subgroups had respective unique cases that exhibited mutually exclusive mutations, suggesting that they may have the redundant effect in the same pathway and selective advantages between them to keep more than one copy of the mutation (**Figure S4**). Other than the mutation pattern, we also investigated differences in the copy number between the two subgroups. GISTIC2.0 software was used to decode the amplification and deletion of CNV on chrmosomes. Compared with C2, C1 had a higher copy number gained in genome and a lower copy number in genome (Figure 6E). The results showed that the two subtypes had frequent copy number variations (CNVs) in the region of oncogenes and tumor suppressor genes (e.g. VHL and TTN), as well as metabolic regulators (e.g. COL9A1 and COL19A1), suggesting that CNVs may play a significant role in the tumorigenesis and progression of ccRCC. The recurrent CNVs in C2 included the amplification of 5q14.3 (NR2F1-AS1), 5q33.2 (KIF4B) and 1p36.11(SYF2), as well as the deletion of 4q24 (PPA2) and 3p21.31 (LTF). The specific CNVs in C1 were mainly associated with cell proliferation, such as the amplification of 5q31.3 (KCTD16), 7p22.2 (SDK1),as well as the deletion of 4q24 (PPA2) (Figure 6F). These results suggest the two subtypes had distinctive CNV events, which not only may cause different immune infiltrations but promote the target treatment of ccRCC.

**Figure 6.**
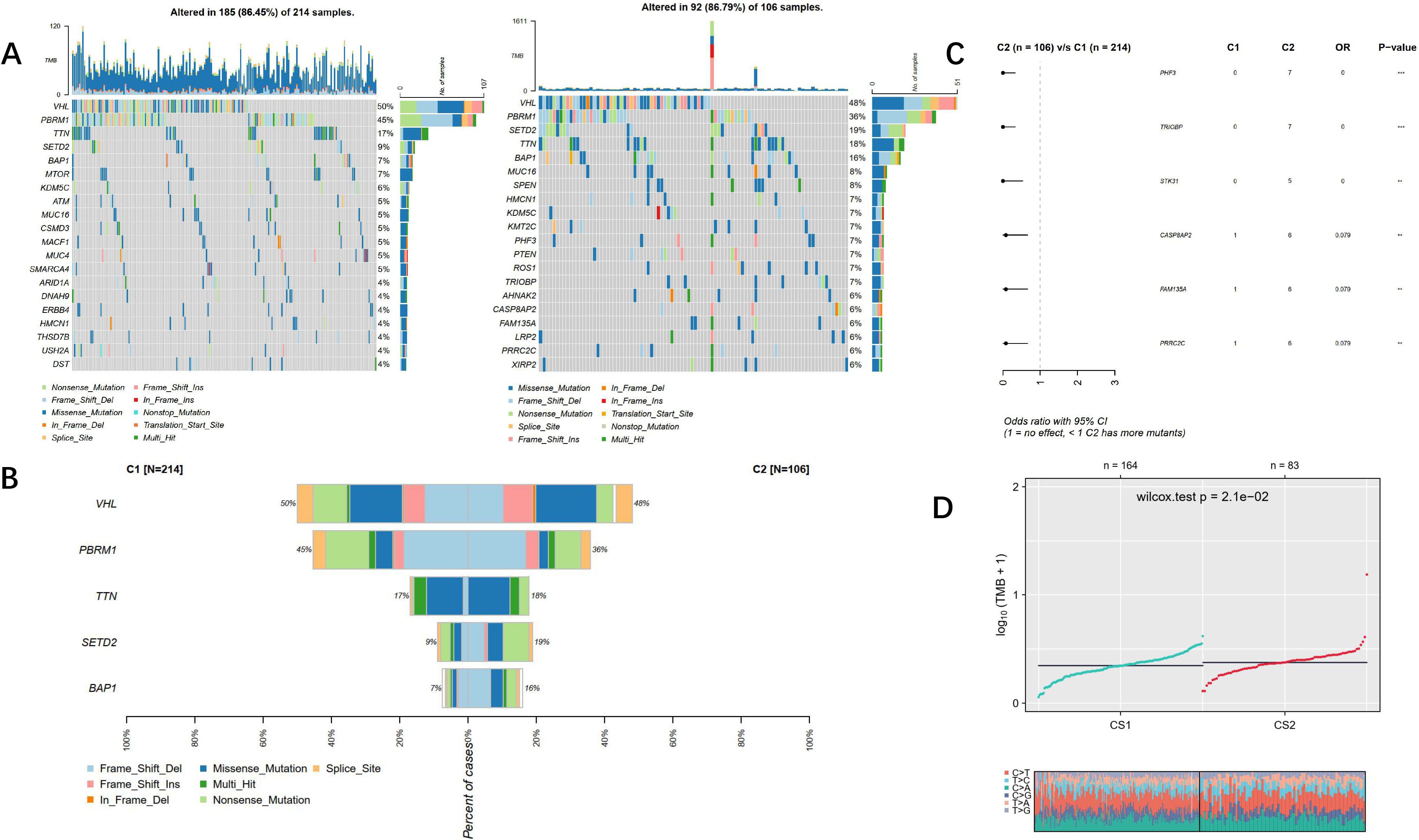
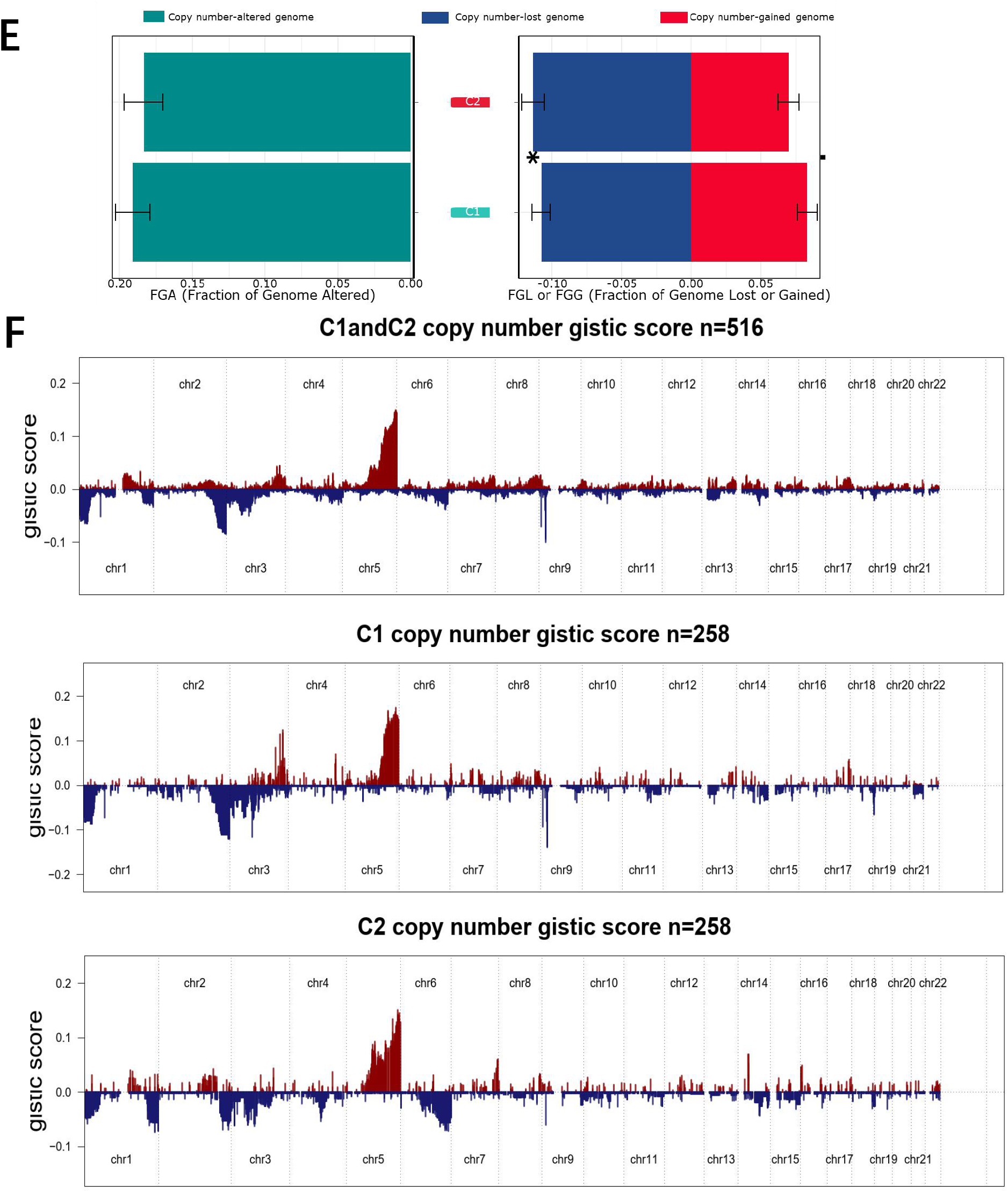
Mutation and CNV differences between subgroups. (A-B) Waterfall plot shows the mutation distribution of the top most frequently mutated genes. (C) Forest plots display the top 6 most significantly differentially mutated genes between the two subgroups. (D) Boxplot of TMB between the two subgroups. (E) Barplot of fraction genome altered in the two identified subtypes. (F) Composite copy number profiles for ccRCC with gains in red and losses in blue and grey highlighting differences.

### Drug sensitivity between the two subgroups

GDSC database was used to forecast the chemotherapy response of the two pyroptosis subtypes to common chemotherapy drugs. It was found that IC50 was significantly different between C1 and C2 subgroups(Figure 7A). At the same time, several potential prodrugs with therapeutic potentialities were investigated in C1 and C2, and the results also showed different IC50values between the two groups (Figure 7B). The detailed molecular structures of these drugs are shown in **Figure S5**.

**Figure 7.**
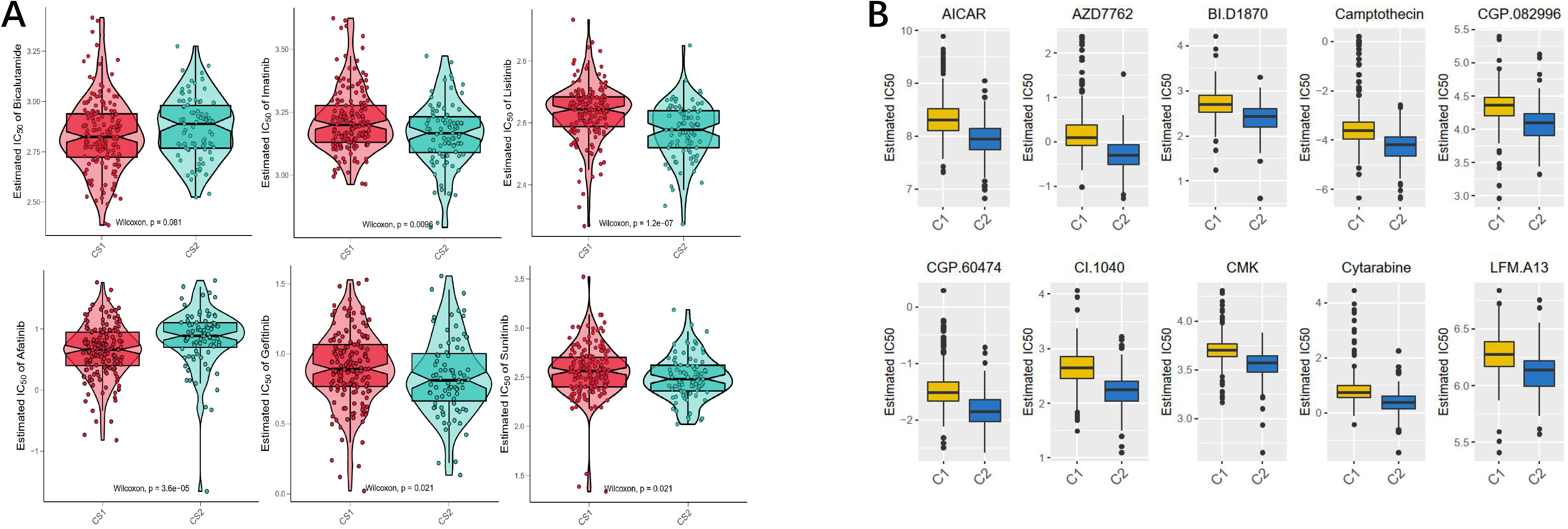
Difference of drug sensitivity. (A) Differences in estimate IC50 of the molecular targeted drugs between the subgroups. (B) The chemotherapy response of the two prognostic subtypes to 10 chemotherapy drugs.

### Construction and validation of a 5-gene pyroptosis related signature model

Firstly, we used univariate analysis to select pyroptosis genes that had impact on OS. Then, the remaining 19 pyroptosis genes were subjected to Lasso-Cox regression analysis and 10-fold cross-validation to generate the optimal model. The Lasso coefficient profile plot was produced against the log(k) sequence, and the minimize k method resulted in 5 optimal coefficients (Figure 8A,B). Finally, a risk model with 5 pyroptosis genes (PYCARD, AIM2, IL6, GSDMB and TIRAP) reached the optimal regression efficiency to speculate the prognostic ability. Furthermore, a pyroptosis risk signature was constructed to estimate the risk score of each patient based on the linear combination of the 5 mRNA expression levels weighted by the Lasso-Cox regression coefficients: Risk score = 0.0271113*AIM2+0.04147645*GSDMB+0.01748664*IL6+0.01968024*PYCARD-0.08678271*TIRA P. To identify the pytoptosis signature responsible for OS and PFI survival prediction, TCGA-ccRCC and ICGC-ccRCC cohort samples were divided into a high-risk group and a low-risk group by using the median risk score as a cutoff point (Figure 8C,D). It was found that OS and PFI in the high-risk TCGA group were poorer than those in the low-risk TCGA group (Figure 8E). To determine whether the risk model had a similar prognostic value in the outer dataset, ICGC-ccRCC cohort was used as a validation dataset. The patients from ICGC dataset were divided to a high-risk group and a low-risk groups based the median risk scores evaluated with the same coefficient. The similar clinical outcome was found in ICGC cohort (Figure 8F). Finally, the different expression level of five risk related genes were verified in SMMU cohort, in which (**Figure S6).**

**Figure 8.**
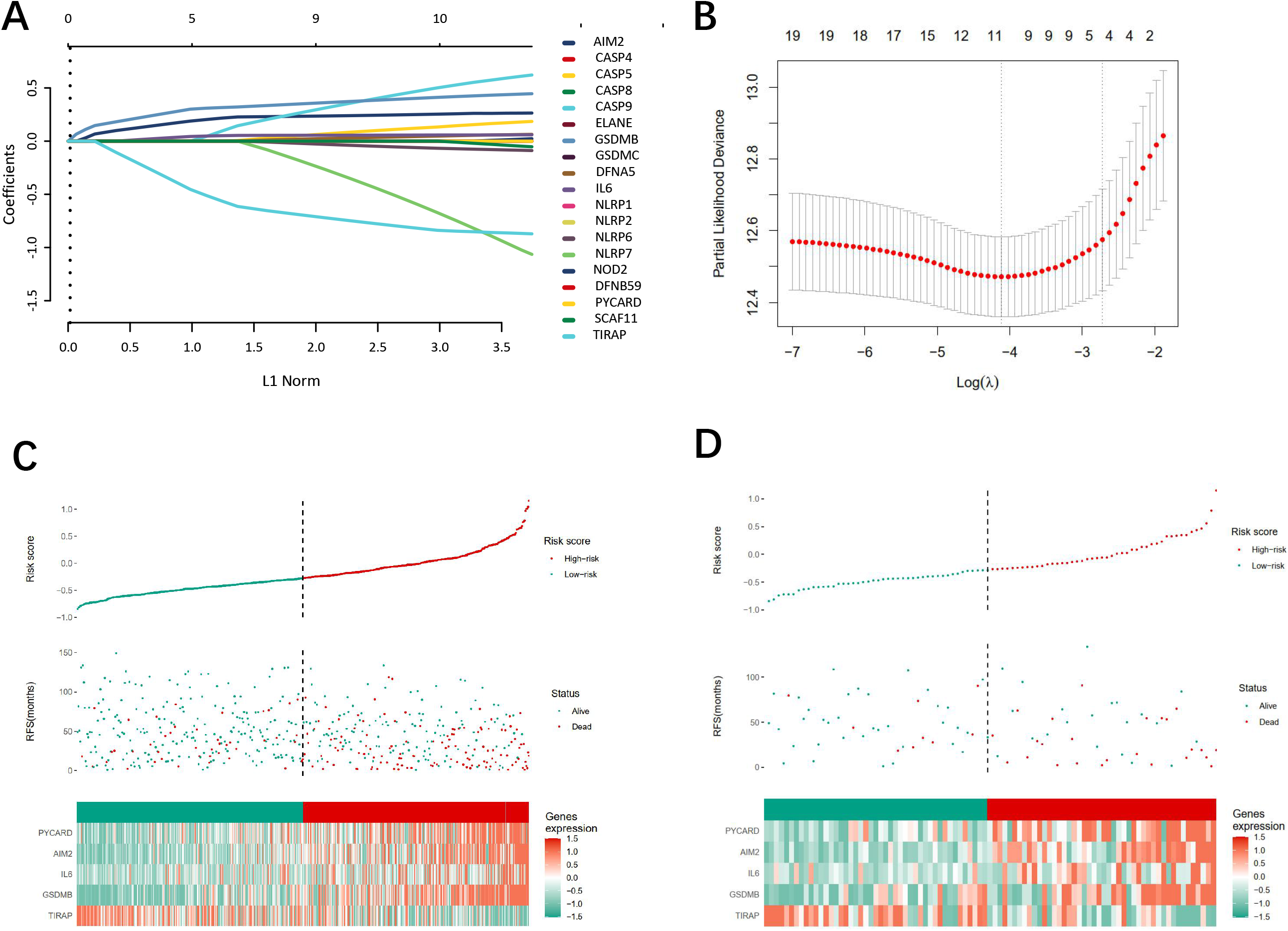
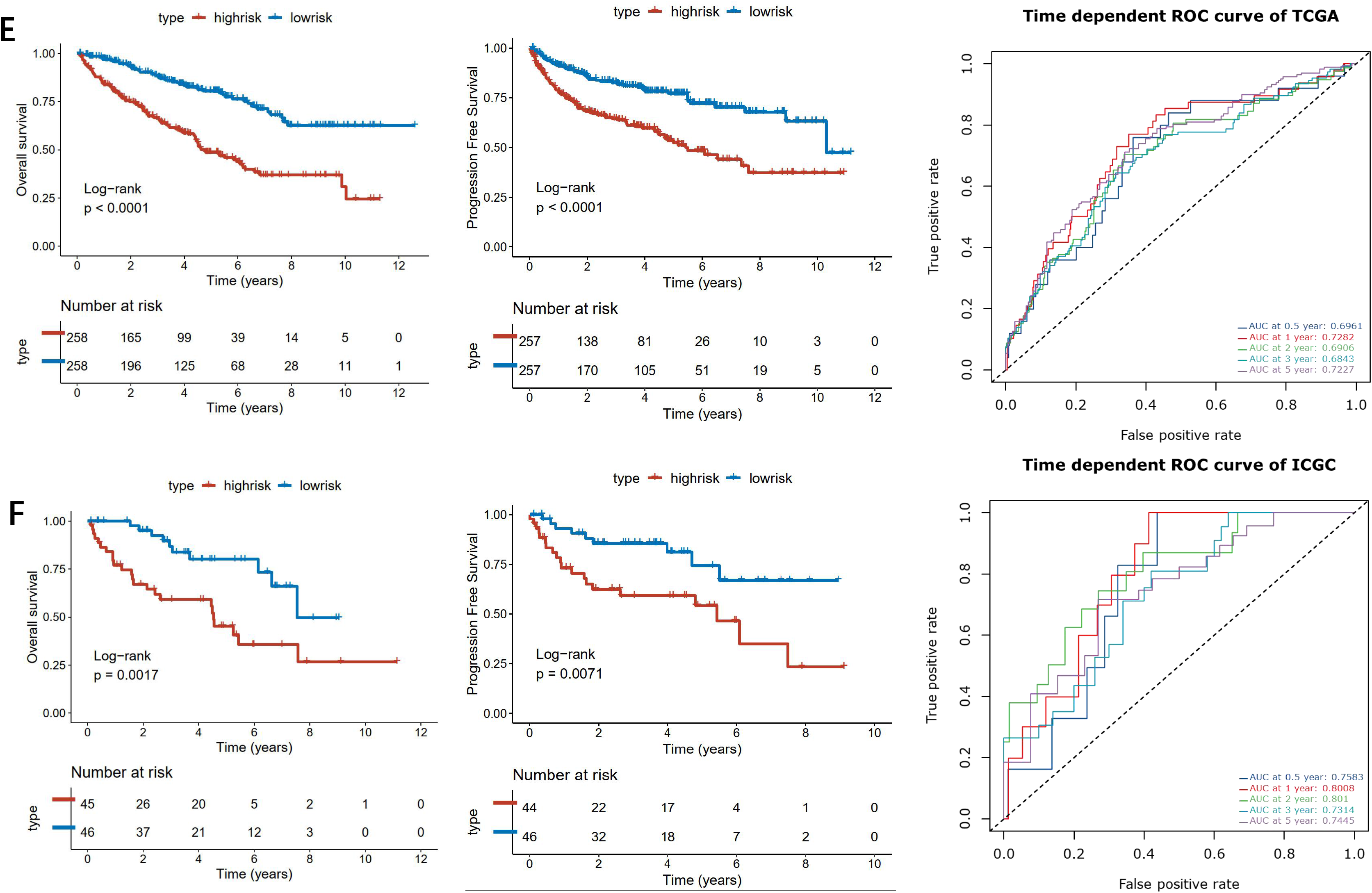
Construction of the pyroptosis related risk scores. (A) LASSO coefficient plot of pyroptosis related genes. (B) The optimal parameter (λ) was chosen by cross validation. (C-D) Risk score analysis in ccRCC patients in the TCGA (left) and ICGC (right) cohort. (E-F) Kaplan-Meier analysis for OS (left) and PFI (right) of the two subtypes in the TCGA and ICGC cohort and the corresponding ROC curve.

## Discussion

Pyroptosis is recognized as a type of programmed cell death and has been demonstrated to be significantly associated with oncogenesis, tumor progression, immune status and anti-tumor response^21^. The role of pyroptosis in urologic carcinoma has attracted increasing attention in recent years. In this study, we firstly analyzed the role of pyroptosis-related genes in ccRCC and found that most of those genes were differentially expressed between ccRCC and normal renal tissues. Then, we identified two heterogeneous pyroptosis-related subgroups (C1 and C2) in ccRCC patients by unsupervised cluster algorithms. It was found that C1 subgroup possessed a high level of pytoptosis-related genes and high abundance of immune cells, which was defined as the pytoptosis ^high^ and immunity ^high^ subtype. C2 expressed low pyroptosis genes and lacked infiltrating immune cells, which was defined as the pyroptosis ^low^ and immunity ^low^ subtype. In addition, we constructed a pyroptosis based risk model based on the pyroptosis signature and evaluated its accuracy and stability in validation datasets, hoping that the results obtained would help better understand the pyroptosis role in ccRCC and promote precise therapy of ccRCC.

It was found in this study that the two subgroups had distinctive clinical characteristics. Patients in C1 subgroup had better OS and PFI relative to C2, and patients in C2 subgroup were associated with worse clinical characteristics in terms of the stage and grade. We further analyzed the drug sensitivity between the two subgroups and found that IC50 in C1 was higher than that in C2 under the treatment of bicalutamide and afatinib, while patients in C2 subgroup were more sensitive to imatinib, lisitinib, gefitinib and sunitinib. In addition, compared with C2, patients in C1 subgroup obtained more benefits from immune therapy as compared those in C2 subgroup.

Next, we investigated genome alterations in C1 and C2 subtypes and found that that SPEN was a particular SMG of C2 and associated with the formation of centrosomes and cilia, and the loss of centrosomes was required for the formation of apoptotic microtubule network. Based on the above evidence, we speculated that SPEN loss in ccRCC may induce pyroptosis formation. SETD2 is a histone H3 K36 methyltransferase that regulates chromatin biology and thereby modulates gene transcription and DNA repair^22^. Intriguingly, SETD2 was recently demonstrated to methylate tubulin for cytoskeleton remodeling and STAT1 for interferon response^23^. It was found in our study that the mutation frequency in C2 subgroup was higher than that in C1. Until now, there has been no research about SETD2 mutation with pytoptosis, and we speculate that SETD2 could enhance pyroptosis in ccRCC.

Additionally, we explored the relationship between ccRCC TME and pyroptosis. Compared with C1, C2 presented more accumulation of immune cell infiltration in TME and higher immune activity. In addition, the expression of the immune check point inhibitor genes was also higher than that in C2. All these findings suggest that the response for immune check point therapy in C1 was significantly higher than that in C2. Given the high pyroptosis status in C2, we speculate that even pyroptosis could not only recruit immune cell infiltration in TME but build an immune suppressive state in TME, suggesting that targeting pyroptosis may convert the immune suppressive state and enhance the immunotherapy effect. Recently, two simultaneously published studies reported that tumor cells undergoing pyroptosis recruit tumor suppressed immune cells, thus waken immune check point inhibitors efficiency^24, 25^.

Based on the pyroptosis status and prognostic characteristics between the two subgroups, we tried to explore and discover different treatment strategies. Interestingly, although TMB in C2 subgroup was higher than that in C1 subgroup, the latter was associated with a better response with ICB therapy, suggesting that the pyroptosis status may play a role in ICB therapy in ccRCC by influencing TME.

Tumorigenesis is a mutagenic process involving participation of the TME component. Recent studies have indicated the necessity of cell-to-cell communications in the progress of various tumors^26^. It is therefore urgent to analyze the role of pyroptosis-related genes in TME. We found that the pyroptosis-related genes were highly expressed in myeloid cells but not in epithelial cells, and T and epithelial cells triggered the pyroptosis effect in other cells in TME via the TNF signaling pathways. Combined with previous immune microenvironment analysis, we speculated that TNF signaling pathways may play an important role in myeloid cells and further mediate the immune impressive environment in ccRCC, and it might reverse the immune impressive state by pyroptosis-medicine target at epithelial cells.

Furthermore, the pyroptosis status may also have impact on sensitivity to chemotherapy. According to the estimate IC50, patients in C1 may be more sensitive to Bicalutamide and Afatinib, while C2 may be sensitize to Imatinib, Lisitinib, Gefitinib and Sunitinib. Based on the pyroptosis status, medical care workers can choose a suitable treatment scheme for patients more accurately. Since the poorer prognosis and lower sensitivity to drug therapy in C2, we used GDSC database to identify small-molecule drugs for C2 patients, including some anti-cancer drugs such as AICAR, Camptothecin and Cytarabine. AICAR (5-Aminotimidazole-4-carboxamide riboside or acadesine) is an AMP-activated protein kinase (AMPK) agonist, which can induce a cytotoxic effect against several cancer cell types. Liang et al found that the combination use of Rapamycin and AICAR could effectively reduce cell proliferation, increase cell apoptosis, and markedly decrease the level of p-Akt, HIF-2α and vascular endothelial growth factor expression in kidney tumor tissues^27^. Camptothecin is a natural anticancer drug in traditional Chinese medicine. Xiao et al found that the Camptothecin analogue G2 could induce apoptosis in liver cancer and colon cancer cell lines by inducing ROC accumulation and reducing MMP^28^. Galley et al reported two types of Camptothecin analogues CPT-11 and 9-AC, which showed a marked survival advantage in an orthotopic model of advanced renal cancer^29^. Cytarabine is an effective drug in the treatment of certain hematologic malignancies. Song et al found that a new generation of cytarabine (Ara-C) analogs could induce the apoptosis effect in prostate cancer via targeting MK2 and inducing the synergistic antitumor activity in p53-deficient prostate cancer cells combining with cabozantinib^30^. Hence, these candidate molecular drugs might also possess potential efficacy for ccRCC.

In this study, we constructed a pyroptosis-related genes risk model and found that it could predict OS and PFI in ccRCC patients both in training and validation cohorts. IL6 is a cytokine that functions in inflammation and maturation of B cells. IL6 is involved in the STATS-mediated signal transduction pathway by mediating tumor immune suppression, tumor cell survival, premetastatic niche formation, and chemotherapy resistance. Yang et al found that CS-Iva-Be, which is a special IL6R antagonist, inhibited IL6/STAT3 signaling pathway and sensitized breast cancer cells to TRAIL-induced cell apoptosis^31^. We found that high IL6 expression was associated with a poor survival outcome in ccRCC, which may be a result of its negative regulation of pyroptosis in ccRCC. GSDMB belongs to the gasdermin (GSDM) family which may adopt different mechanisms of intramolecular domain interactions to modulate their lipid-binding and pore-forming activities. The GSDM family has regulatory functions in cell proliferation and differentiation, especially in the pyroptosis process. Previous studies on human cancers have demonstrated that GSDMB is highly expressed in both healthy and cancer tissues including gastric, uterine, cervical and breast cancers^32^. Researchers have demonstrated that GSDMB is located in the amplicons, genomic regions that are often amplified during cancer development. Therefore, GSDMB may be involved in cancer progression and metastasis. In this study, we found that GSDMB was highly expressed in ccRCC, and this high expression was positively correlated with ccRCC progression. TIRAP is an important adaptor protein belonging to the TLR/IL-1R superfamily, with a TIR domain in the cytoplasmic tail. It was reported that aberrant expression of TIRAP could induce the development of multiple tumors including lymphocytic leukemia, gastric cancer and colorectal cancer^33^. It was also found that phycocyanin, a food derived inhibitor, could inhibit TIRAP in NSCLS cells and exert an anti-proliferation effect through down-regulating TIRAP/NF-kB activity in lung cancer^33^. We found that phycocyanin could also serve as a protective factor in renal cancer patients, and its expression was positively correlated with the renal cancer stage. AIM2 is a protein of the interferon-inducible PYRIN and HIN domain-containing (PYHIN) family. Some recent studies reported that AIM2 acted as a DNA sensor in innate immunity by directly binding to foreign double-stranded DNA in infected macrophages^34^. AIM2 could trigger the assembly of inflammasomes to induce a caspase1 mediated inflammatory response, causing cell apoptosis. Chen et al found that exogenous AIM2 expression could reduce breast cancer cell proliferation by inhibiting the nuclear factor kappa-B (NF-κB) transcriptional activity and suppressing mammary tumor growth^35^. PYCARD is a pro-apoptotic gene encoding a signaling factor that consists of an N-terminal PYRIN-PAAS-DAPIN domain (PYD) and a C-terminal caspase-recruitment domain (CARD) and operates in the intrinsic and extrinsic cell death pathways. Miao et al found that a lncRNA antisense to PYCARD exhibited a dual nuclear and cytoplasmic distribution and promoted proliferation of cancer cell lines by downregulating the expression of PYCARD^36^. Several studies found that PYCARD was a tumor-inhibiting factor in that it was silenced in many tumor types and the level of methylation in its promoter was negatively correlated with tumor progression^37^. We found that PYCARD was highly expressed in tumor cells and its expression was positively correlated with renal tumor progression. Kumari et al found that exogenous overexpression of AIM2 could suppress the tumorigenicity in immunocompromised nude mice by suppressing mTOR-S6K1 pathways^38^. Our study found that AIM2 was highly expressed in renal tumor tissues as compared with that in normal renal tissue, and its high expression was positively correlated with poor prognosis of ccRCC patients.

Although Ours study has provided a more comprehensive view into the role of pyroptosis in ccRCC and established a powerful model for prognostic prediction, there are still two major drawbacks that require further exploration. Firstly, we only analyzed the prognostic role of the pyroposis-related genes in ccRCC, but how these genes interact with each other during pyroptosis remains to be further investigated. Secondly, although we performed an independent internal validation, it is difficult to cover all variations in patients from different geographical regions when tissues and information were retrospectively collected in publicly available databases.

## DECLARATIONS

### Ethics approval and consent to participate

As the data used in this study are publicly available, no ethical approval was required.

### Consent for publication

Not applicable

### Availability of data and material

The raw data for this study were generated at the corresponding archives. Derived data supporting the findings are available from the corresponding author [WLH] on reasonable request.

### Competing interests

The authors have no conflict of interest.

### Funding

This study was funded by the National Natural Science Foundation of China (82072812, 81730073, 81872074), Shanghai Sailing Program (19YF1448300) and Clinical science and technology innovation project of Shanghai Shenkang Hospital Development Center (SHDC12018108).

## Acknowledgements

We greatly appreciate the patients and investigators who participated in the corresponding medical project for providing data. We thank Dr.Jianming Zeng(University of Macau), and all the members of his bioinformatics team, biotrainee, for generously sharing their experience and codes.

---

**Table S1** Primer sequences of 5 hub genes

**Figure S1.** Workflow of this study

**Figure S2.** (A)Enrichment analysis of DEGs; (B)Transcription factor analysis of dysregulated genes; the purple nodes represent down-regulated genes(left) and un-regulated genes(right), and green nodes represent transcription factors.

**Figure S3.** (A) Heatmap showing the selected ligand-receptor interactions between cancer cells and other cells via CellChat; (B) Bubble plot showing the selected ligand-receptor interactions between cancer cells and other cells via cellphonedb.

**Figure S4.** Heatmap illustrates the mutually co-occurring and exclusive mutations of the top 25 frequently mutated genes. The color and symbol in each cell represent the statistical significance of the exclusivity or co-occurrence for each pair of genes.

**Figure S5.** The structure tomographs of the ten candidated small-molecule drugs.

**Figure S6.** Different expression level of five risk related genes in SMMU cohort.

## Notes

https://www.cancer.gov/about-nci/organization/ccg/research/structural-genomics/tcga

https://dcc.icgc.org/

### Competing Interest Statement

The authors have declared no competing interest.

