## Supplementary figures and images for "Establishment of a prognosis prediction model based on pyroptosis-related signatures associated with the immune microenvironment and molecular heterogeneity in clear cell renal carcinoma"

### Supplementary Figures

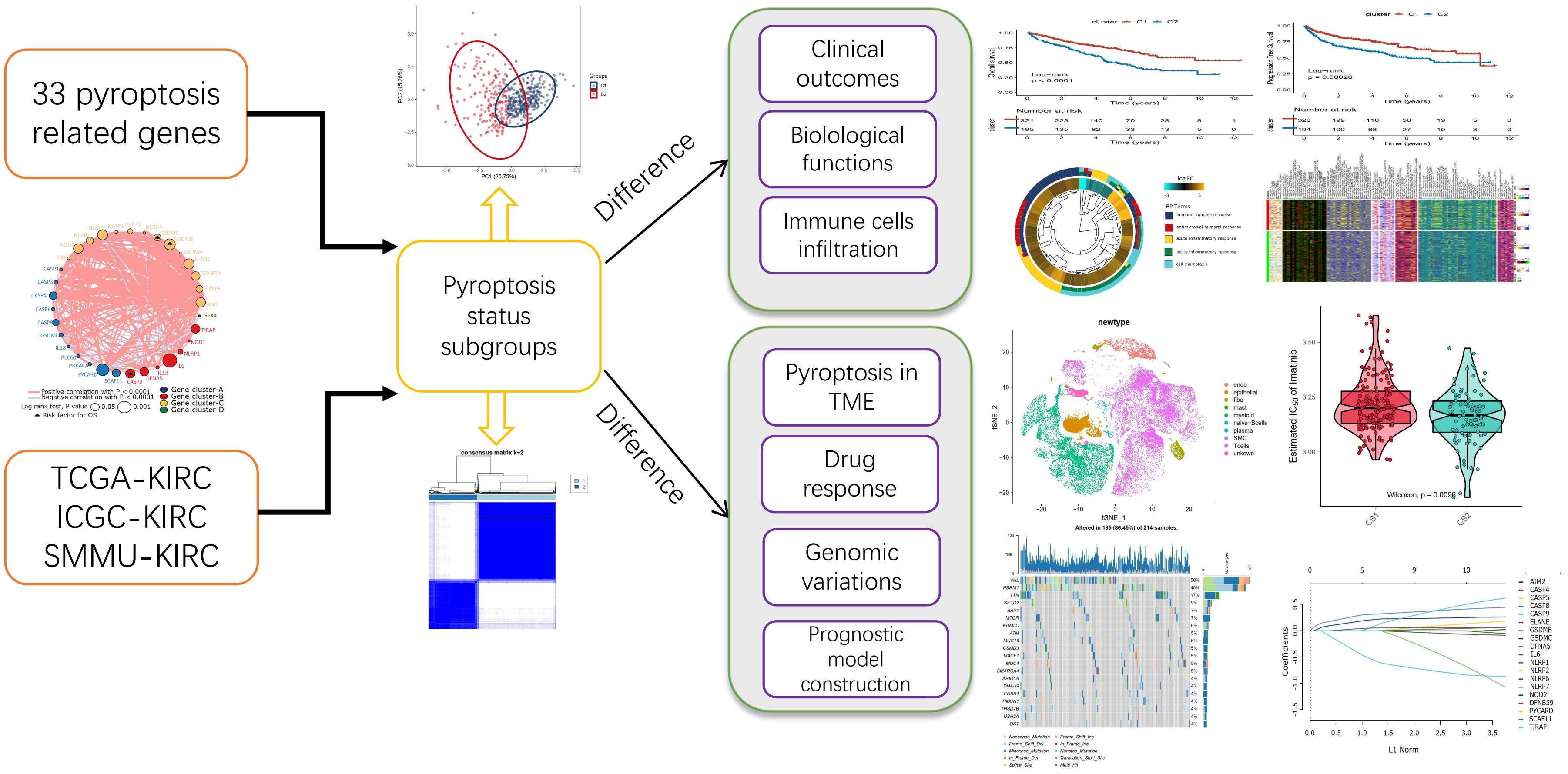

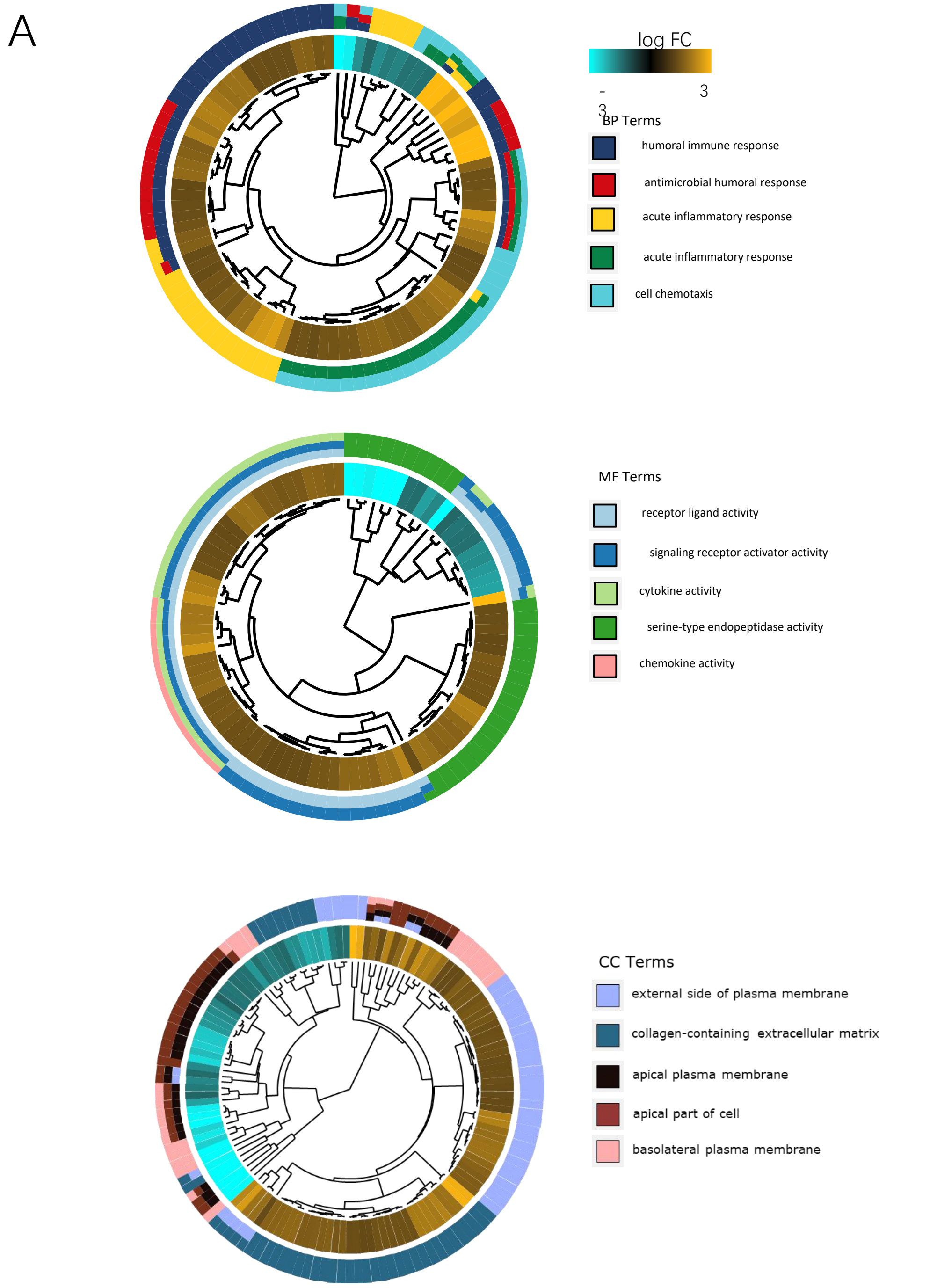

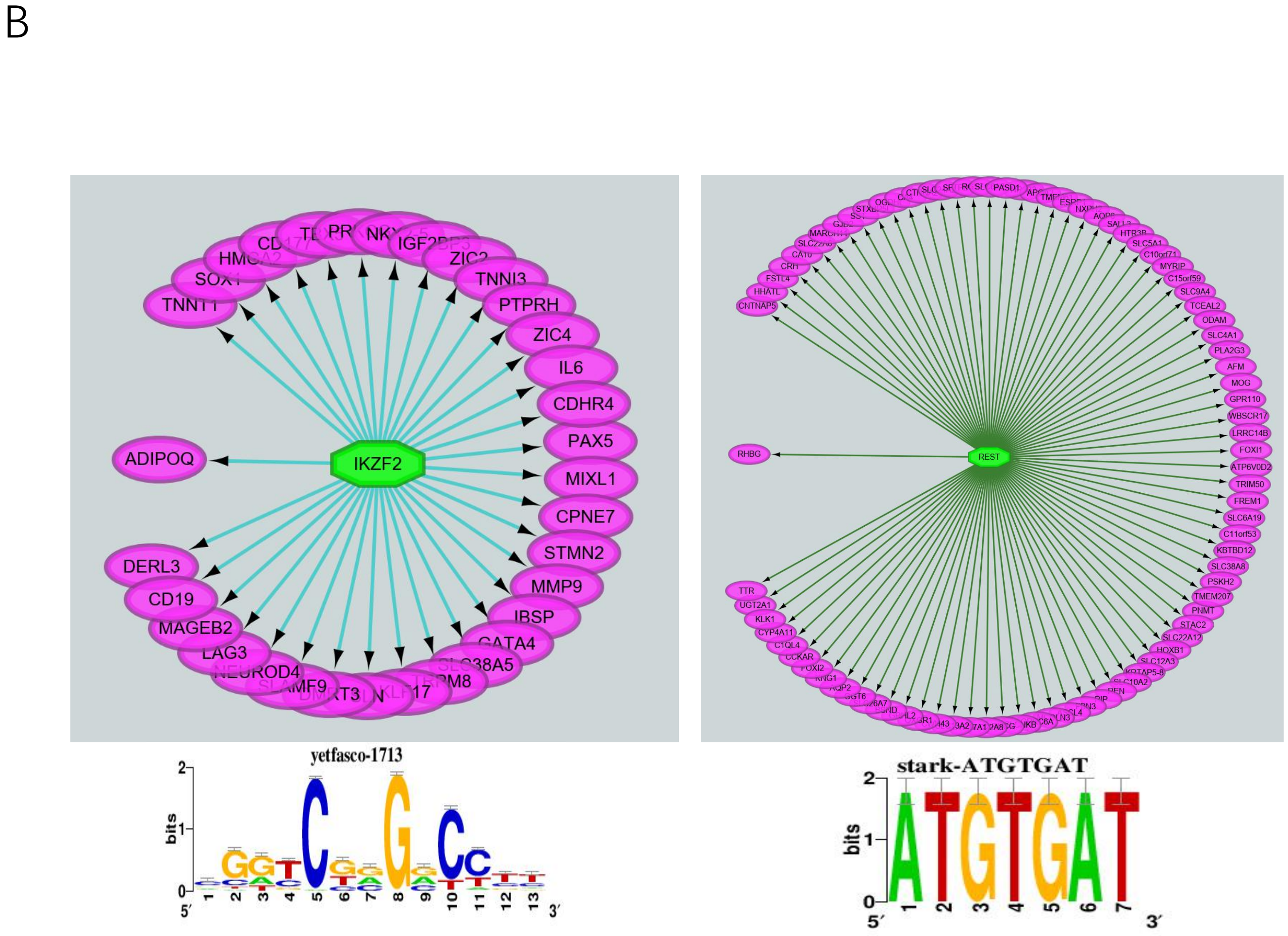

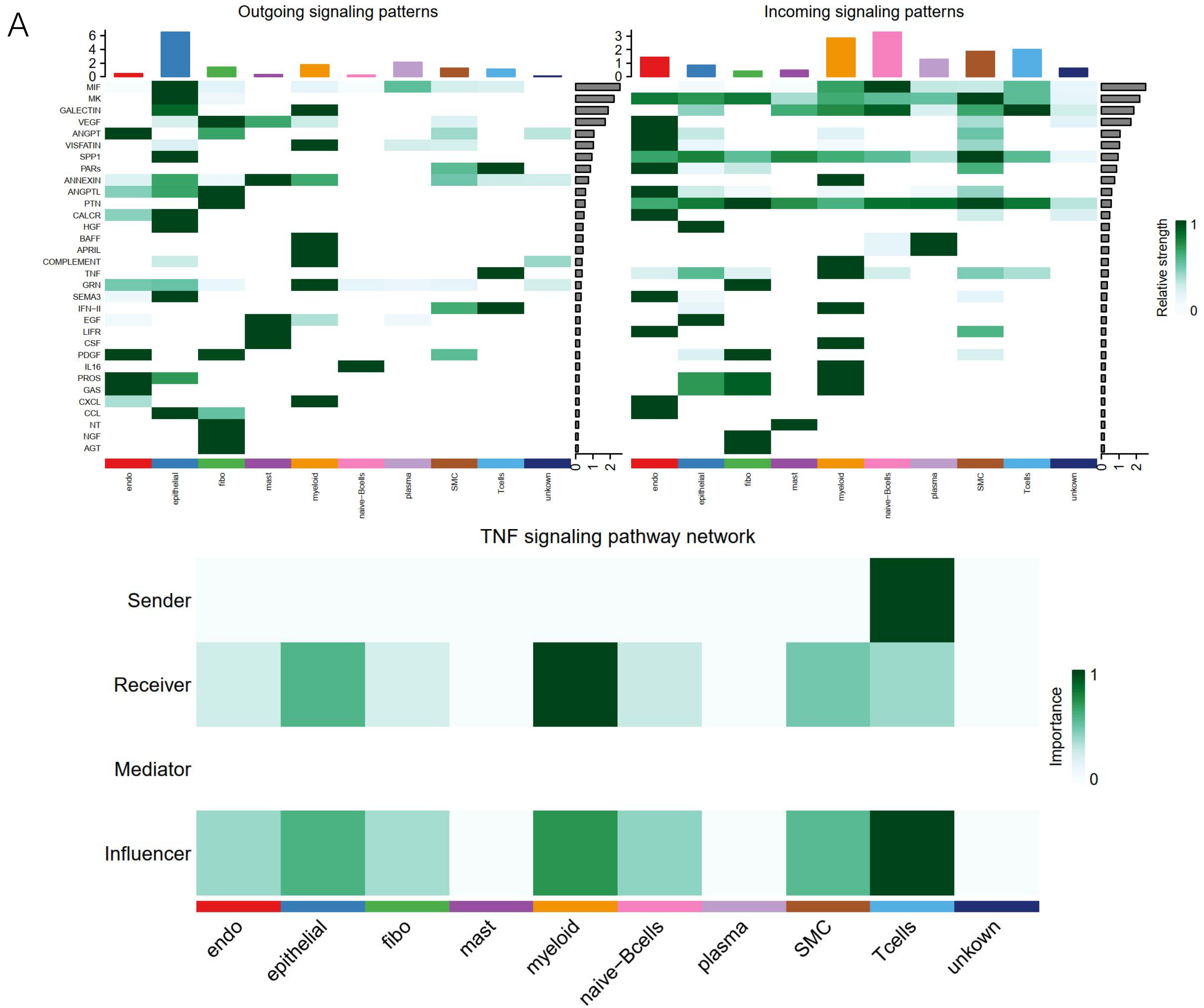

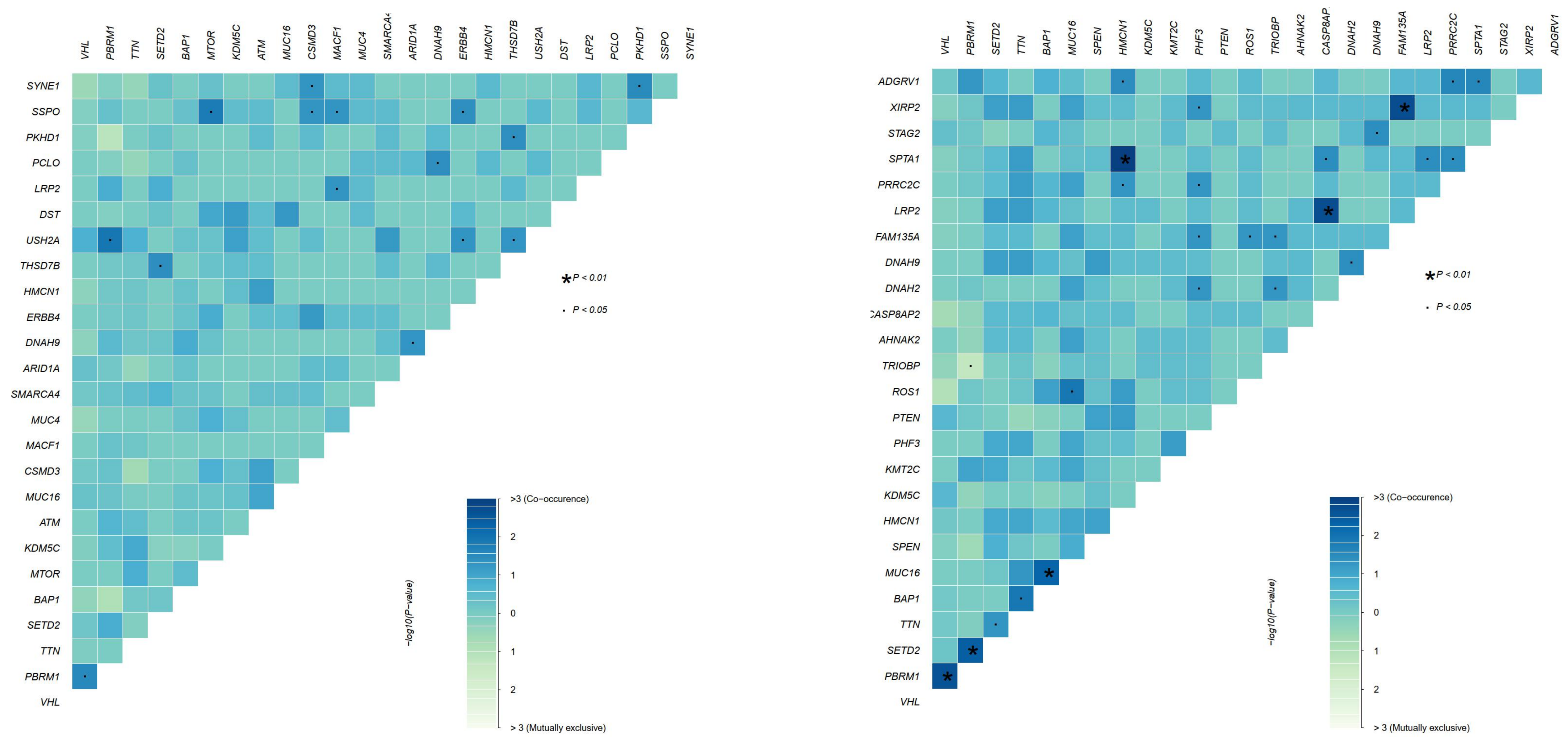

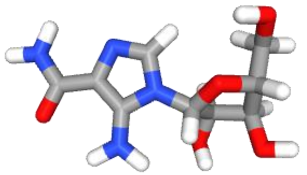

AICAR

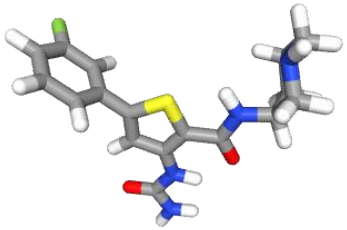

AZD7762

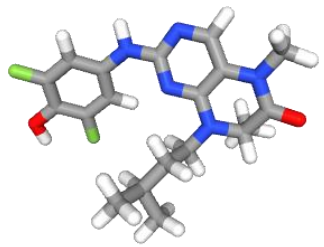

BI.D1870

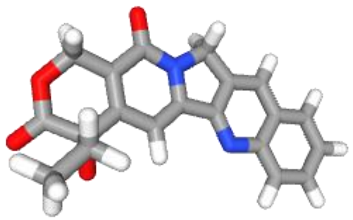

Camptothecin

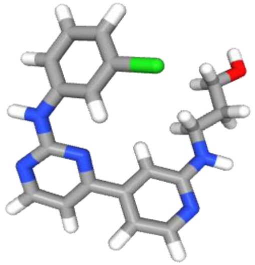

CGP.082996

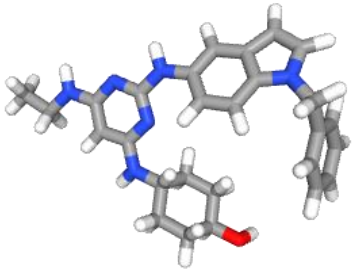

CGP.60474

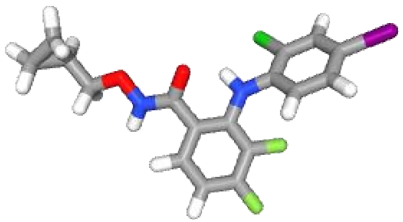

CI.1040

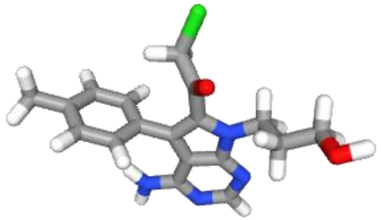

CMK

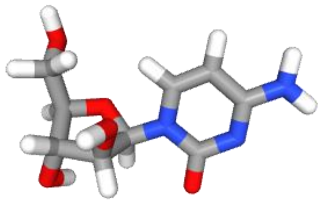

Cytarabine

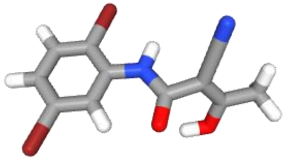

LFM.A13

Figure S5

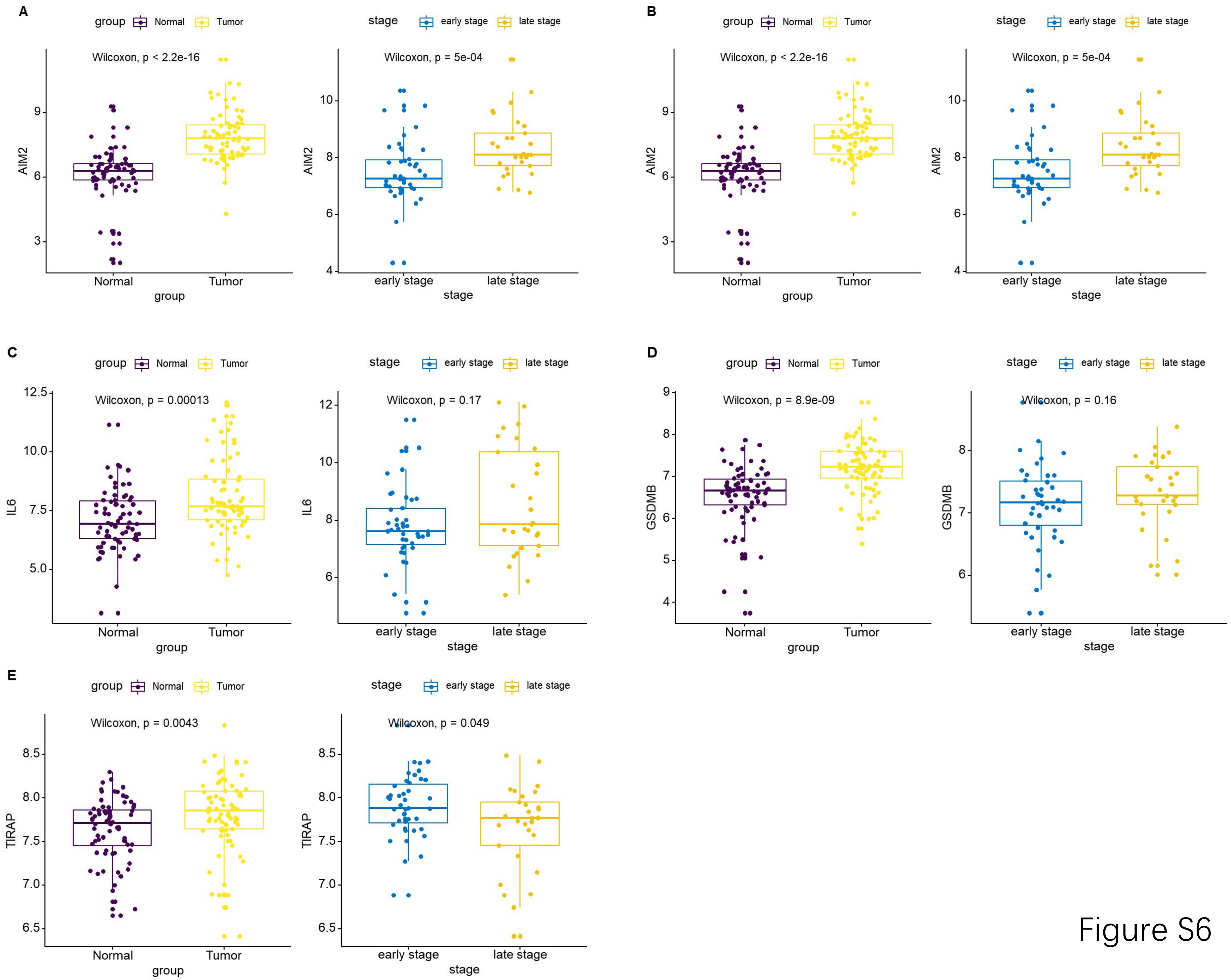
