## Supplementary Table for "Establishment of a prognosis prediction model based on pyroptosis-related signatures associated with the immune microenvironment and molecular heterogeneity in clear cell renal carcinoma"

PYCARD

F: TGGATGCTCTGTACGGGAAG

R: CCAGGCTGGTGTGAAACTGAA

AIM2

F: TGGCAAAACGTCTTCAGGAGG

R: AGCTTGACTTAGTGGCTTTGG

IL6

F: ACTCACCTCTTCAGAACGAATTG

R: CCATCTTTGGAAGGTTCAGGTTG

GSDMB

F: TGATTGCCGTTAGAAGCCTTG

R: TCCCGTTGAGTCTACATTATCCA

TIRAP

F: ATGGCATCATCGACCTCCCT

R: GTCACTCGCATGTGTGGGT
